# Genomic Structural Equation Modeling Reveals Latent Phenotypes in the Human Cortex with Distinct Genetic Architecture

**DOI:** 10.1101/2022.11.04.515213

**Authors:** Rajendra A. Morey, Yuanchao Zheng, Delin Sun, Melanie E. Garrett, Marianna Gasperi, Adam X. Maihofer, Lexi Baird, Katrina L. Grasby, Ashley Huggins, Courtney C. Haswell, C. Paul M. Thompson, Sarah Medland, Daniel E. Gustavson, Matthew S. Panizzon, William S. Kremen, Caroline M. Nievergelt, Allison E. Ashley-Koch, Mark W. Logue

**Author notes:** <u>Corresponding Author</u>: Rajendra A. Morey MD, Associate Professor of Psychiatry and Behavioral Sciences, Faculty Member of the Duke Institute for Brain Sciences, Member of the Center for Brain Imaging and Analysis, VISN 6 MIRECC, Durham VA Health Care System, Durham, NC, USA. 3022 Croasdaile Drive, Suite 300, Durham NC 27705, USA.

## Abstract

Genetic contributions to human cortical structure manifest pervasive pleiotropy. This pleiotropy may be harnessed to identify unique genetically-informed parcellations of the cortex that are neurobiologically distinct from anatomical, functional, cytoarchitectural, or other cortical parcellation schemes. We investigated genetic pleiotropy by applying genomic structural equation modeling (SEM) to model the genetic architecture of cortical surface area (SA) and cortical thickness (CT) of 34 brain regions recently reported in the ENIGMA cortical GWAS. Genomic SEM uses the empirical genetic covariance estimated from GWAS summary statistics with LD score regression (LDSC) to discover factors underlying genetic covariance. Genomic SEM can fit a multivariate GWAS from summary statistics, which can subsequently be used for LD score regression (LDSC). We found the best-fitting model of cortical SA was explained by 6 latent factors and CT was explained by 4 latent factors. The multivariate GWAS of these latent factors identified 74 genome-wide significant (GWS) loci (p<5×10^−8^), including many previously implicated in neuroimaging phenotypes, behavioral traits, and psychiatric conditions. LDSC of latent factor GWAS results found that SA-derived factors had a positive genetic correlation with bipolar disorder (BPD), and major depressive disorder (MDD), and a negative genetic correlation with attention deficit hyperactivity disorder (ADHD), MDD, and insomnia, while CT factors displayed a negative genetic correlation with alcohol dependence. Jointly modeling the genetic architecture of complex traits and investigating multivariate genetic links across phenotypes offers a new vantage point for mapping genetically informed cortical networks.

**HIGHLIGHTS:**

- Genomic SEM can examine genetic correlation across cortical regions.
- We inferred regional genetic networks of cortical thickness and surface area.
- Network-associated variants have been implicated in multiple traits.
- These networks are genetically correlated with several psychiatric disorders including MDD, bipolar, ADHD, and alcohol dependence.

## 1. INTRODUCTION

A number of different neurobiological markers have been employed in conjunction with various organizational schemes to map the human cortex. It is possible that individual differences in regional cortical surface area (SA) and cortical thickness (CT) and may drive factors that affect each person and each region independently. However, the covariance structure of regional SA and CT reveals that individual differences are systematically coordinated within communities of brain regions, fluctuate in magnitude together within a population, may be instantiated as structural covariance networks (SCN)^1^, and partially recapitulate established organizational schemes^2–5^. For instance, SCN organization is consistent with topological patterns of cortical maturation observed throughout developmental stages from childhood and adolescence into early adulthood ^6^, and the same patterns are then targeted by neurodegenerative diseases in late life^7,8^. Second, brain regions with highly correlated CT or SA often represent networks that perform dedicated cognitive processes^1,9,10^. Third, regions within SCNs tend to be directly connected by white matter tracts. Indeed, about 40% of SCN connections show convergent white matter fiber connections, although other relationships captured by SCNs are independent of fiber connectivity^5^.

The correlation structure between regions represented by an SCN is influenced by both the environment and genetics. The genetic factors underlying structural correlations closely resemble functional and developmental patterns^4,5,11^. We will refer to these patterns of genetic correlations between brain regions as *genetically informed brain networks* (GIBNs). Genetic correlations of CT or SA have been examined with twin studies^12,13^. These genetic influences were recapitulated in over 400 twin pairs, to show that the cortex is organized genetically into communities of structural and functional regions, is hierarchical, is modular, and is bilaterally symmetric^11^. Their genetically informed parcellation identified 12 spatially contiguous regions that qualify as GIBNs. Relatedly, SA and CT phenotypes overlap genetically with GIBNs, the latter being less granular and more discoverable^14^.

While twin studies have laid important groundwork regarding genetic correlations of the brain, they have several limitations. First, twin studies do not provide specific genetic variants associated with each GIBN^11^ and therefore offer an incomplete characterization of cortical pleiotropy. Second, twin studies rely on the *equal environment* assumption, which may by invalid for some studies. Third, quantifying the genetic correlation between GIBNs and low prevalence traits such as schizophrenia (0.5% prevalence)^15^ or bipolar disorder (1% prevalence)^16^ would require an extraordinarily large number of twin pairs to wield sufficient statistical power. Recently, genetic correlations between brain regions derived from GWAS results have been applied to estimate the contribution of common genetic variation. This method confers several advantages over twin studies. These SA and CT GWAS results reveal pleiotropy and genetic correlation across many neuroimaging phenotypes^17,18^. Additionally, genome-wide SNP data allow effect-size estimation for individual variants and the ability to test genetic correlations with other traits in different populations.

Genomic structural equation modeling (gSEM) is a multivariate statistical method that leverages the genetic architecture of <u>many</u> genetically correlated phenotypes to derive relatively <u>few</u> latent phenotypes, which explain the observed genetic correlation and loadings of multiple phenotypes onto a latent phenotype^19^. Therefore, gSEM applied to GWAS offers a genetically informed parcellation of the cortex that is neurobiologically distinct from anatomical, functional, cytoarchitectural, and other parcellation schemes^6,20^. Latent factors represent traits that explain the genetic correlation across multiple regions and define the brain regions that constitute each GIBN. Importantly, gSEM estimates the strength of association between genetic variants and each latent factor followed by a multivariate GWAS of each GIBN using GWAS summary statistics for individual correlated traits. Importantly, gSEM provides a description of the underlying genetic architecture of the traits being examined and effect size estimates for the underlying latent factors.

In the present study, we sought to elucidate the genetic architecture of 34 regional SA and CT phenotypes reported in the ENIGMA-3 GWAS of over 50,000 primarily healthy individuals. We hypothesized that gSEM might identify brain partitions consistent with the 12 clusters described by Chen et al.^11^, along with other viable solutions. The genetic correlations reported in Grasby et al.^18^, were stronger within major anatomical lobes than across lobes. Thus, while we predicted gross lobar structure may be reflected by GIBNs, we further predicted that GIBNs would reflect the complex relationships captured by functional networks, canonical resting-state networks, fiber tract networks, gene expression networks, and other neurobiological systems^6,11^. We hypothesized from the outset that most genetic variants discovered by the ENIGMA-3 cortical GWAS would influence GIBNs in the present study, but we also sought to discover novel genetic markers, and discover links between known genetic variants and established regional associations as well as GIBNs. Additionally, we hypothesized genetic correlations between GIBNs and major neuropsychiatric disorders.

## 2. METHODS

### 2.1 Data

We used the results of the ENIGMA-3 cortical GWAS that identified genetic loci associated with variation in cortical SA and CT measures in 51,665 individuals primarily (∼94%) of European descent, from 60 international cohorts^18^. Phenotype measures were extracted from structural MRI scans for 34 regions defined by the Desikan-Killiany atlas using gyral anatomy, which establishes coarse partitions of the cortex^21^. Two global measures of total cortical SA and average CT, as well as 34 regional measures of SA and CT were averaged across left and right hemisphere structures to yield 70 distinct phenotypes. Multiple testing correction in the ENIGMA-3 GWAS was based on 60 independent phenotypes with a GWS threshold of P≤8.3×10^−10^. We accessed the GWAS summary results for the 34-regional bilateral analyses performed by Grasby et al. The primary GWAS presented in Grasby et al. had adjusted for global measures (total SA and mean CT). However, we requested alternate results without global adjustments to avoid artefactual negative (inverse) correlations between regions. Regional SA and CT metrics were analyzed separately.

### 2.2 Analysis

Our analyses were performed using the genomic-SEM package which is available for the R programming language^19^. The entire gSEM was performed twice, once for 34 SA regions and once for 34 CT regions. Like standard SEM, gSEM includes an exploratory factor analysis (EFA) stage and a confirmatory factor analysis (CFA) stage. To avoid overfitting between the confirmatory and exploratory phases, we analyzed odd chromosomes in the EFA and even chromosomes in the CFA. Whereas SEM often fits multiple models corresponding to *a priori* hypotheses built on theoretical models, we took a hypothesis free (data driven) approach. In the EFA, we fit one model, which included up to 10 factors, for each of SA and CT. The optimal number of factors for each was determined using scree plots (see Supplementary **Figures S1** and **S2**). Positive factor loading estimates greater than a pre-specified threshold were carried forward to the confirmatory factor model stage to be re-estimated, and the remaining loading parameters were set to zero. As there was no consensus on factor loading cutoffs ^22,23^, we tested thresholds 0.3 and 0.5. Cross loadings that were allowed if they exceeded the threshold. Factors that loaded on only a single region were removed from CFA modeling. Therefore, some models with a large number of factors ended up as redundant, and were not carried forward to CFA.

All of the distinct factor structures generated were carried forward to CFA and re-estimated using even chromosomes. Standardized root-mean square residual (SRMR), Akaike Information Criteria (AIC), model □^2^, and Comparative Fit Index (CFI) were used to evaluate model fit of the CFA models. The large number of GWASs and the large sample size of each GWAS meant that all model □^2^ statistics were highly significant (p∼0) and hence are not presented.

The top performing factor models in the CFA were further optimized by successive removal of non-significant factor loadings. In addition, we attempted to fit a bifactor model as part of the CFA step to account for correlation between the factors. Specifically, we fit a bifactor model where a “total” CT or SA factor was added, which loaded on all regions, and a multi-level model where all factors from the EFA loaded onto a 2^nd^ order factor. The bifactor model failed to converge for all CFA models and the multilevel model failed to improve model fit; hence these results are not presented. For our purposes, a model which minimized the AIC was deemed optimal. SRMR and CFI were calculated to measure model fit.

Once the GIBNs were defined, we used gSEM to generate a multivariate GWAS of each GIBN. The GWS associations (p<5×10^−8^) for each GIBN were compared to the significant SNPs reported by Grasby et al. (with global correction) and then compared to the corresponding GWAS results without the global correction (the same results used to generate GIBNs). The <u>FU</u>nctional <u>M</u>apping and <u>A</u>nnotations (FUMA) package^24^ was used to annotate results from each GIBN GWAS, including annotating SNPs to specific genes, and identifying potential functional variants. FUMA was run based on LD in the 1000G Phase3 EUR reference panel^25^ and the default FUMA parameter settings.

### 2.3 Polygenicity Analysis

We examined the significant SNPs from the GIBN GWAS, as well as SNPs in LD using FUMA to test for functional associations with established behavioral traits and major neuropsychiatric disorders. First, we examined whether observed variants from the GWAS recapitulated GWS SNPs from previous GWAS results of neuroimaging traits including cortical GWAS results and other structural neuroimaging parameters^17,26–31^. We also looked for SNPs that were significant in GWASs of 12 neuropsychiatric disorders from the Psychiatric Genomics Consortium (PGC): ADHD^32^, alcohol dependence^33^, anorexia nervosa^34^, autism spectrum disorder^35^, bipolar^36^, cannabis use^37^, MDD^38^, obsessive compulsive disorder (OCD)^39^ posttraumatic stress disorder (PTSD)^40^, schizophrenia^41^, Tourette”s syndrome^42^, and anxiety^43^. Finally, FUMA was used to functionally annotate loci that overlapped with previously published GWAS results.

#### Genetic Correlation with Psychopathology

We used cross-trait LDSC to identify links between psychiatric disorders and CT-derived GIBNs as well as psychiatric disorders and SA-derived GIBNs^44^. We estimated the genetic correlation between CT- and SA-derived GIBNs and neuropsychiatric disorders using their GWAS summary statistics^44^. To maximize statistical power, we limited the number of genetic correlations to 12 neuropsychiatric disorders.

### 2.4 Data and Code Availability

The GWAS summary statistics which were used in this paper are available to download from the ENIGMA consortium website (http://enigma.ini.usc.edu/research/download-enigma-gwas-results). Access to cohort data is available either through public repositories or directly from the cohort. Direct requests are required when informed consent or the approved study protocol does not permit deposition into a repository. Requests for data by qualified investigators are subject to scientific and ethical review to ensure that the data will be used for valid scientific research and to ensure compliance with confidentiality, consent, and data protection regulations. Some of the data are subject to material transfer agreements or data transfer agreements, and specific details on how to access data for each cohort are available in Grasby et al (2020).The Genomic SEM package used to analyze the data is publicly available at https://github.com/GenomicSEM/GenomicSEM. The ldsc package is publically available at https://github.com/bulik/ldsc. The results of the multivariate GWASs of the CT- and SA-derived GIBNs are available at https://pgc-ptsd.com/about/workgroups/imaging-workgroup/.

## 3. RESULTS

### 3.1 Model Fit

The SA-derived 6-GIBN solution resulted in the best overall model fit to the genetic covariances generated from the GWAS summary statistics (AIC=22,712,604, CFI=0.920, SRMR=0.062). See Supplementary **Table S1** for fit statistics for each evaluated model. The 6 SA-derived GIBNs (SA1-SA6) loaded on 24 of the 34 brain regions^18^. The standardized estimates for the 6 SA-derived GIBN models (standardized factor loadings) are presented in Supplementary **Table S2** and presented in **Figure 1**. The GIBNs generally encompass contiguous brain regions and many correspond to known neuroanatomical features. SA1 contains loadings for inferior temporal, isthmus cingulate, postcentral, precuneus, superior parietal, supramarginal, and temporal pole. SA2 contains loadings for caudal anterior cingulate, caudal middle frontal, medial orbitofrontal, paracentral, and rostral anterior cingulate. SA3 contains loadings for banks superior temporal sulcus (STS), inferior parietal, and middle temporal. SA4 contains loadings for pars opercularis, pars orbitalis, and pars triangularis, SA5 contains loadings for cuneus, lateral occipital, lingual, and pericalcarine, and SA6 corresponds to the auditory cortex. The 6-factor model indicated substantial correlation between GIBNs (r_g_=0.61 to 0.91) as reported in Supplementary **Table S3**.

**Figure 1.**
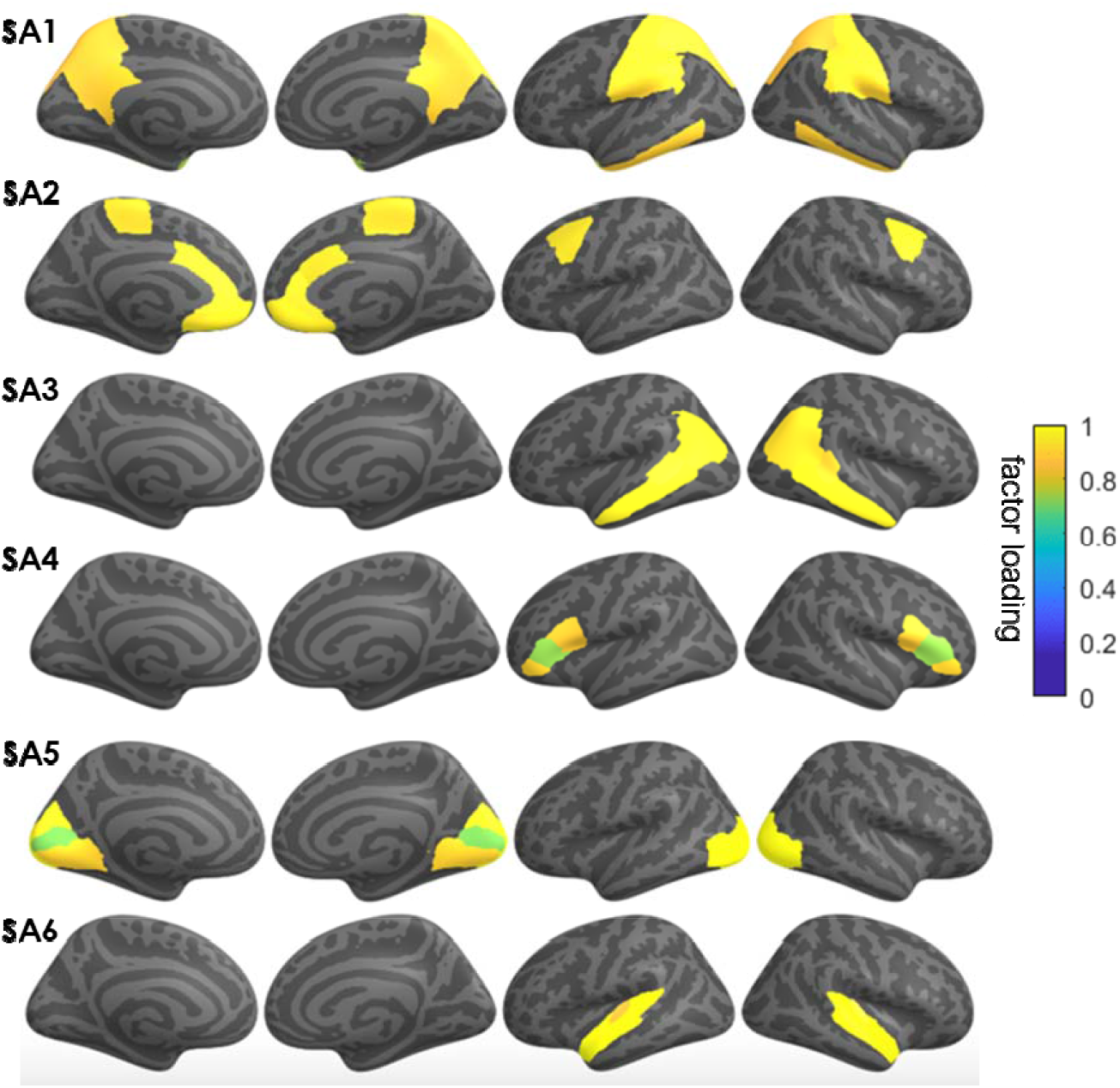
Genomic structural equation modeling (gSEM) jointly modeled the genetic architecture of cortical surface area (SA) for 34 brain regions based on GWAS results of Grasby et al (2020). The model generated 6 *genetically Informed brain networks* (GIBNs) from SA phenotype measures. The color overlay on cortical regions represents the magnitude of the factor loadings indicated in the color gradient (yellow = high; blue = low). Subsequent GWAS identified several genome wide significant hits (p < 5×10^−8^) associated with each GIBN.

The CT-derived 4-GIBN solution resulted in the best model fit (AIC=17761928, CFI=0.932, SRMR=0.077; Supplementary **Table S4**). Significant non-zero loadings for CT-derived GIBNS loaded on 25 of the 34 brain regions from Grasby et al. See Supplementary **Table S5** for the estimated loadings that are visualized in **Figure 2**. As observed with SA models, the CT-derived GIBNs generally encompassed contiguous cortical regions. CT1 contains loadings for banks STS, caudal middle frontal, inferior parietal, paracentral, pars opercularis, post-central, pre-central, precuneus, rostral middle frontal, superior frontal, superior parietal, and supramarginal cortices. CT2 contains loadings for caudal anterior cingulate, frontal pole, insula, lateral orbitofrontal, medial orbitofrontal, pars orbitalis, rostral anterior cingulate, and rostral middle frontal. CT3 contains loadings for banks STS, superior temporal, and temporal pole. CT4 contains loadings for cuneus, lateral occipital, parahippocampal, and pericalcarine cortices. The CT-derived GIBNs were moderately to highly correlated (r_g_=0.67 to 0.87; **Supplementary Table S6**).

**Figure 2.**
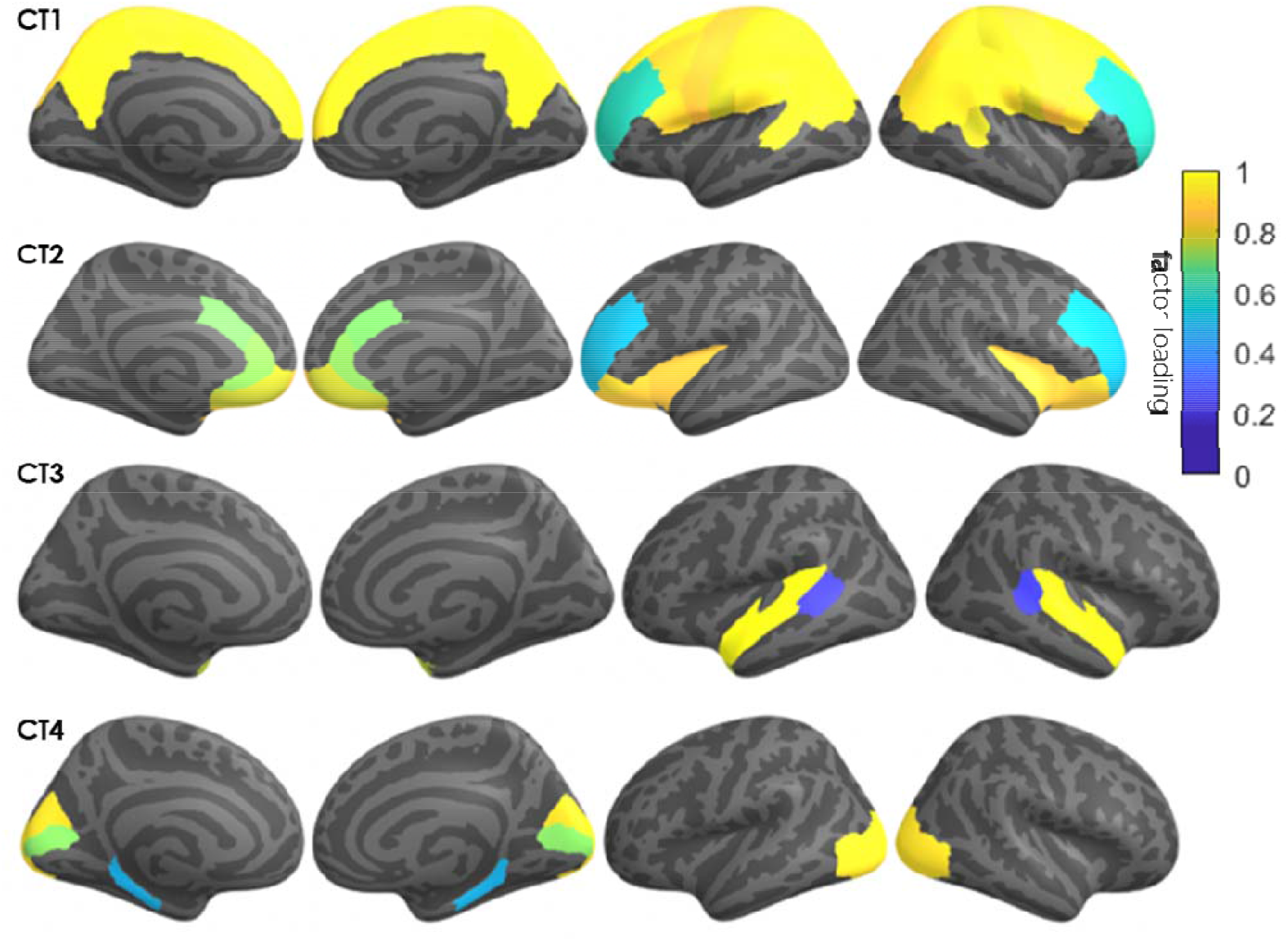
Genomic structural equation modeling jointly modeled the genetic architecture of cortical thickness (CT) for 34 brain regions based on GWAS results of Grasby et al (2020). The model generated 4 *genetically informed brain networks* (GIBNs) from CT phenotype measures. The color overlay on cortical regions represents the magnitude of the factor loadings indicated in the color gradient (yellow = high; blue = low). Subsequent GWAS identified several genome wide significant hits (p < 5×10^−8^) associated with each GIBN.

Factor diagrams for SA- and CT-derived GIBNs are presented in **Figure 3**. Consistent with prior work, the SA-derived GIBNs were largely distinct from CT-derived GIBNs, although some regional overlap exists between SA- and CT-derived GIBNs. For example, SA5 and CT4 are both 4-region GIBNs, with 3 overlapping regions.

**Figure 3.**
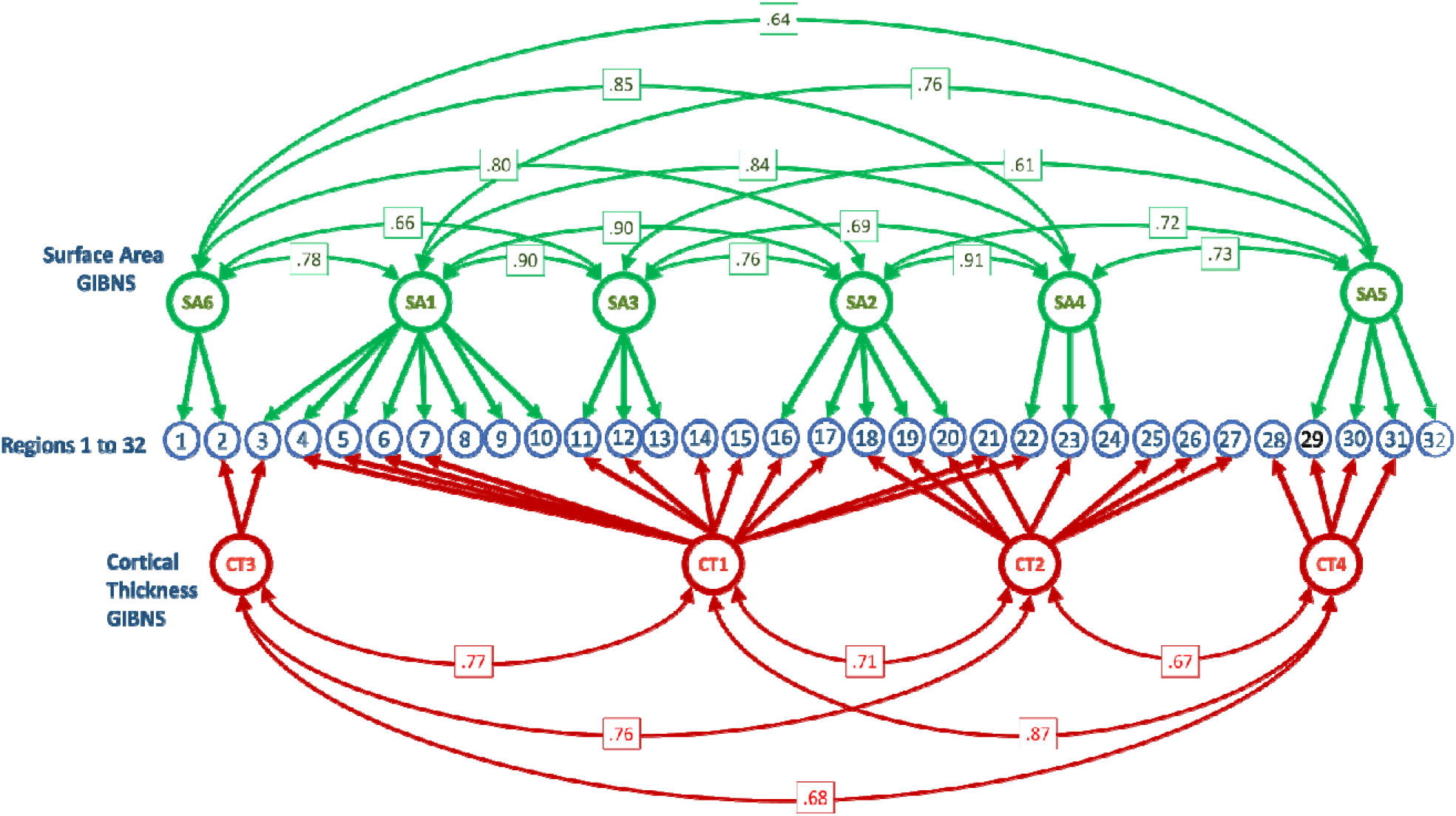
Graph of genomic structural equation modeling (gSEM) results. The blue circles, numbered from 1 to 32, represent the cortical surface area (SA) and cortical thickness (CT) of regions defined by the Desikan-Killiany atlas. Latent SA variables, indicated by green circles, represent the genetic contributions from regional SA, which are specified by thick green lines and arrows. Latent CT variables, indicated by red circles, represent the genetic contributions from regional SA, which are specified by thick red lines and arrows. Thin green lines connect genetically latent SA variables with their genetic correlation strength (r_0_) indicated in green boxes. Thin red lines connect genetically latent SA variables with their genetic correlation strength (r_0_) indicated in red boxes.

### 3.2 GWAS of Genetically Informed Brain Networks

To identify specific genetic variants that may be influencing the GIBNs, we performed a multivariate GWAS on each SA- and CT-derived GIBN. Manhattan plots for SA- and CT-derived GIBN GWASs, their associated quantile-quantile (QQ) plots, and genomic inflation factors (λ) are provided in **Figures S3** to **S12**. We observed moderate p-value inflation (λ values between 1.06 and 1.16). However, the single-trait LD Score regression intercepts for SA- and CT-derived GIBNs were all less than 1.02, indicating that the apparent inflation was likely due to pleiotropy. A total of 5,843 GWS (p<5×10^−8^) variants were associated with 10 GIBNs. Annotation by FUMA ^24^ mapped these variants to 74 independent regions, including 64 loci that were associated with the 6 SA-derived GIBNs and 10 loci that were associated with the 4 CT-derived GIBNs. A phenogram^45^ of the 74 genetic associations is presented in **Figure 4**. A list of all GWS loci with their associated GIBNs is provided in Supplementary **Table S7** Except for two novel SNPs, all others were previously identified in Grasby et al.^18^ in either the analysis adjusted for global SA/CT or the unadjusted analysis. The first novel SNP rs3006933, which resides near the genes *SDCCAG8* and *AKT3* on chromosome 1, was associated with SA1 (p=4.08×10^−9^). The other novel SNP rs1004763 on chromosome 22, which resides in the vicinity of gene *SLC16A8*, was associated with CT2 (p=3.41×10^−08^).

**Figure 4.**
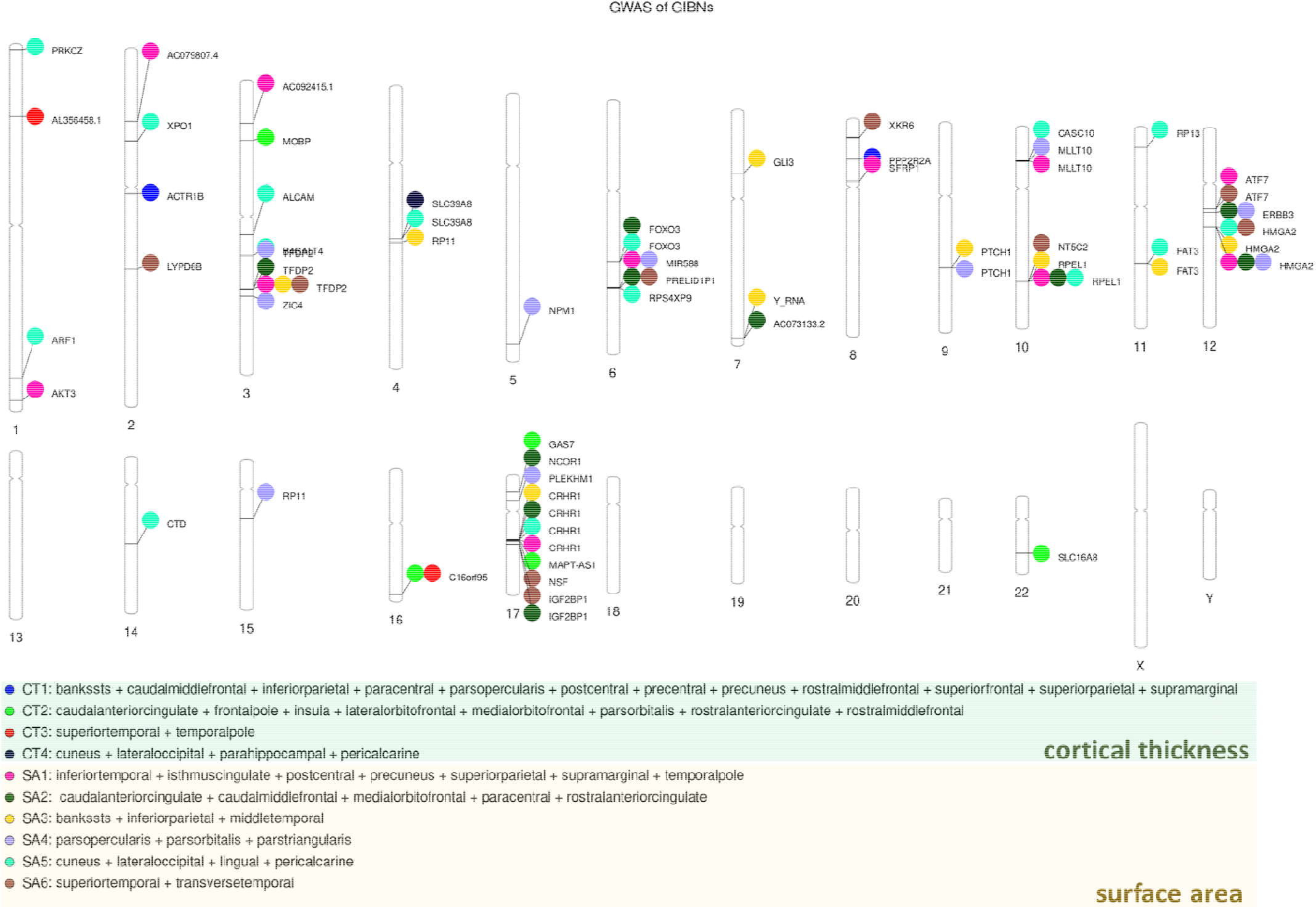
Phenogram of GWS SNPS associated with six genetically informed brain networks (GIBNs) derived from surface area (SA; gold inset) and four GIBNs derived from cortical thickness (CT; green inset).

### 3.3 Genetic Correlation

We observed significant genetic correlation between multiple traits and SA-derived GIBNs as reported in Supplementary **Table S8**. ADHD exhibited significant negative genetic correlation with all SA-derived GIBNs except SA4 (r_g_=−0.13 to −0.20, p=3.29×10^−6^ to 0.0038, p_FDR_=0.00040 to 0.039). Significant positive genetic correlations were observed between bipolar disorder and SA1, SA2, SA4, and SA5 (r_g_=0.10 to 0.14, p=3.00×10^−4^ to 0.0047, p_FDR_=0.012 to 0.043). Interestingly, we observed significant genetic correlations between MDD and SA-derived GIBNs, but in the opposite direction as bipolar disorder. We found a significant negative correlation between MDD and SA6, which was not associated with bipolar disorder (r_g_=−0.10, p=0.0011, p_FDR_=0.17). Negative nominally significant (uncorrected p<0.05) correlations were observed between MDD SA1-SA3, and SA5 (r_g_= −0.057 to −0.080, p=0.0090 to 0.046), while SA4 was not genetically correlated with MDD (p=0.12). SA4 was significantly correlated with cannabis use disorder (r_g_=0.15; p=4.00 × 10^−4^, p_FDR_=0.012), while SA2 was nominally associated with cannabis use (r_g_=0.11, p=0.011). No significant genetic correlations were observed between the 6 SA-derived or 4-CT derived GIBNs and anorexia, autism, anxiety, schizophrenia, PTSD, or Tourette”s Syndrome (all p>0.05).

Fewer genetic correlations were significant between CT-derived GIBN regions and psychiatric disorders (Supplementary **Table S9**). CT3 and CT4 were negatively correlated with alcohol use disorder, exhibiting the strongest correlations with any traits that we examined (CT3 r_g_=−0.35, p=3×10^−4^, p_FDR_=0.012; CT4 r_g_=−0.31, p=7×10^−4^, p_FDR_=0.014). We found a negative nominally significant correlation between alcohol use disorder and CT1 (r_g_=−0.18, p=0.035, p_FDR_=0.22). CT3 had a positive nominally significant correlation with OCD (r_g_=0.22, p=0.0091, p_FDR_=0.078).

The overlap of the GIBN GWS loci with prior GWAS of neuroimaging phenotypes or psychiatric disorders firmly points to the relevance of GIBN-related variants to brain structure and cognition. First, we note that novel variant rs3006933 has been previously associated with subcortical volumes^46^. Novel variants rs3006933 and rs1004763^17,29^ have been associated with neuroimaging phenotypes of corpus callosum white matter microstructure^47^. A comparison of our GIBN GWAS with published psychiatric disorder GWAS results found that multiple SNPs linked to SA-derived GIBNs were also implicated in a GWAS of schizophrenia^41^. Specifically, we identified a cluster of 4 loci in the *CRHR1* gene strongly associated with SA-derived GIBNs (rs62057153 associated with SA1) in our GWAS (p=5.22×10^−17^ to 8.45×10^−21^). We also observed an association between CT1 and rs11692435 (p=1.17×10^−12^), a schizophrenia-related locus, within the *ACTR1B* gene. Finally, CT- and SA-derived GIBNs were associated with schizophrenia risk variants in the *SLC39A8* gene; namely rs13107325 was associated with CT5 and rs13135092 was associated with SA5. No other traits had GWS variants associated with any of the GIBNs.

Annotation of GIBN-related SNPs using FUMA found that many have been previously associated with cognitive, behavioral, neuroanatomical, neurofunctional, and neuropsychiatric phenotypes. In addition to rs3006933 noted above^14^, SA6-linked locus rs9909861^48^ and SA5-linked SNP rs7570830^14^ have also been associated with subcortical volumes. Multiple loci associated with SA-derived GIBNs that encompass temporal, parietal, and temporo-parietal association cortices include SA1-linked locus rs10109434^49^, SA3-linked SNP rs2299148^50^, and SA6-linked locus rs9909861^50–54^ have previously been implicated in academic attainment and cognitive ability. These GIBNS, particularly temporal (SA6) and temporoparietal (SA3) cortices, are the most more strongly linked to academic attainment and the most heritable^55^. The SA5-linked locus rs6701689 has been reported for risk tolerance^56^. However, there is no support for this GIBN in the visual cortex (SA5) plays a role in risk tolerance, which is linked to cerebellar, midbrain, and prefrontal cortical anatomy, as well as glutamatergic and GABAergic neurotransmission^56,57^. The CT4-associated locus rs13107325 has been associated with many traits including schizophrenia^58–65^, bipolar disorder^62,63^, Parkinson”s disease^64,65^, sedentary behavior^46,66^ and risk taking^56^, as well as cognition, intelligence, and educational attainment^50–54,67^. This GIBN includes the parahippocampal and fusiform gyri, which have a firmly established link to schizophrenia^68^ and a recently identified link to sedentary behavior^69^.

## 4. DISCUSSION

The goal of the present study was to leverage the pleiotropic architecture of the human cortex to construct a genetically informed parcellation that could be distinct from anatomical, functional, cytoarchitectural, or other established parcellation schemes, although, our analysis starts with a somewhat crude anatomically-based 34-region parcellation. We investigated the genetic pleiotropy of regional cortical morphology by applying gSEM to jointly model the genetic architecture of 34 brain regions using results from the ENIGMA-3 GWAS^18^. The process was undertaken with gSEM to generate several possible solutions, from which the best-model fit was selected. This solution organized brain regions to optimally assign genetic pleiotropy to 6 SA- and 4 CT-derived latent factors, which we have termed *genetically informed brain networks* (GIBNs). Subsequent multivariate GWAS of these GIBNs were mapped by FUMA to 74 independent SNPs (p<5×10^−8^). LDSC results of CT- and SA-derived GIBNs were positively correlated with OCD and bipolar disorder, but negatively correlated with alcohol use disorder, ADHD, MDD, and insomnia.

We observed 74 GWS markers associated with the SA- and CT-GIBNs. The overlap of the GIBN GWS loci with prior GWAS of neuroimaging phenotypes or psychiatric disorders firmly points to the relevance of GIBN-related variants to brain structure and cognition. First, we note that novel variant rs3006933 has been previously associated with subcortical volumes^46^. Novel variants rs3006933 and rs1004763^17,29^ have been associated with neuroimaging phenotypes of corpus callosum white matter microstructure^47^. A comparison of our GIBN GWAS with published psychiatric disorder GWAS results found that multiple SNPs linked to SA-derived GIBNs were also implicated in a GWAS of schizophrenia^41^. Specifically, we identified a cluster of 4 loci in the *CRHR1* gene strongly associated with SA-derived GIBNs (rs62057153 associated with SA1) in our GWAS (p=5.22×10^−17^ to 8.45×10^−21^). We also observed an association between CT1 and rs11692435 (p=1.17×10^−12^), a schizophrenia-related locus, within the *ACTR1B* gene. Finally, CT- and SA-derived GIBNs were associated with schizophrenia risk variants in the *SLC39A8* gene; namely rs13107325 was associated with CT5 and rs13135092 was associated with SA5. No other traits had GWS variants associated with any of the GIBNs.

Annotation of GIBN-related SNPs using FUMA found that many have been previously associated with cognitive, behavioral, neuroanatomical, neurofunctional, and neuropsychiatric phenotypes. In addition to rs3006933 noted above^14^, SA6-linked locus rs9909861^48^ and SA5-linked SNP rs7570830^14^ have also been associated with subcortical volumes. Multiple loci associated with SA-derived GIBNs that encompass temporal, parietal, and temporo-parietal association cortices include SA1-linked locus rs10109434^49^, SA3-linked SNP rs2299148^50^, and SA6-linked locus rs9909861^50–54^ have previously been implicated in academic attainment and cognitive ability. These GIBNS, particularly temporal (SA6) and temporoparietal (SA3) cortices, are the most more strongly linked to academic attainment and the most heritable^55^. The SA5-linked locus rs6701689 has been reported for risk tolerance^56^. However, there is no support for this GIBN in the visual cortex (SA5) plays a role in risk tolerance, which is linked to cerebellar, midbrain, and prefrontal cortical anatomy, as well as glutamatergic and GABAergic neurotransmission^56,57^. The CT4-associated locus rs13107325 has been associated with many traits including schizophrenia^58–65^, bipolar disorder^62,63^, Parkinson”s disease^64,65^, sedentary behavior^46,66^ and risk taking^56^, as well as cognition, intelligence, and educational attainment^50–54,67^. This GIBN includes the parahippocampal and fusiform gyri, which have a firmly established link to schizophrenia^68^ and a recently identified link to sedentary behavior^69^.

The GIBNs we generated can be compared to similar structures generated from twin studies. Using 400 twin pairs, Chen et al. generated twelve genetically-informed clusters from vertex-based surface area measures^11^. The 12 clusters consisted of (1) motor-premotor cortex, (2) dorsolateral prefrontal cortex, (3) dorsomedial frontal cortex, (4) orbitofrontal cortex, (5) pars opercularis and subcentral region, (6) superior temporal cortex, (7) posterolateral temporal cortex, (8) anteromedial temporal cortex, (9) inferior parietal cortex, (10) superior parietal cortex, (11) precuneus, and (12) occipital cortex. The results of Chen et al.^11^ constitute the earliest genetically informed parcellation of the brain, which reported heritability estimates and genetic correlations between clusters. The genetically informed clusters are more consistent with classical anatomically-defined sulcal and gyral boundaries, Brodmann definitions, and cytoarchitectural patterns than our GIBNs. Importantly, our GIBNs, better represent functional specializations than the 12 genetically-informed clusters. However, these published twin studies notably lack any information about specific genetic variants. Thus, our study extends the circumscribed results of earlier twin studies by mapping specific genetic variants onto the cortical covariance structure.

The organization of structural covariance networks is partially reflected in other schemes for organizing the human cortex including resting state networks, gene expression networks, white matter networks and other neurobiological brain features. Structural covariance networks (SCN) tend to reflect white matter connections throughout the cerebral cortex, although other information captured by SCNs is independent of fiber connectivity^5^. More recently, the covariance structure of cortical thickness was shown to be correlated with gene transcriptional networks that are organized with similar complex topology on the basis of modularity, small-worldness, and rich-clubness ^2^. Whereas network properties such as degree and degree-distribution differed, the cortical areas connected to each other within SCN modules had higher levels of gene-co-expression than expected by chance alone^2^. Relatedly, gene co-expression networks are also mirrored by canonical resting-state networks^70^. Specifically, 136 consensus genes are differentially co-expressed in 4 resting-state networks including salience, dorsal default mode, visual, and sensorimotor.

It is now firmly established that resting-state functional connectivity from fMRI evinces robust patterns of synchronous activity that intrinsically organize into canonical resting-state networks ^71^. These resting state networks are typically identified by applying independent component analysis to the functional connectome^72^. We found that SA-derived GIBNs significantly align with several canonical resting-state networks. Most prominent among them is the recapitulation of the visual network by SA5, which is composed of cuneus, lateral occipital, lingual, and pericalcarine cortices^73^. Twin-based non-linear multidimensional heritability estimates (of multidimensional traits such as brain network architecture) are among the highest for the visual network (left h^2^_m_=0.53; right h^2^_m_=0.45) and auditory network (left h^2^_m_=0.44; right h^2^_m_=0.60)^74^. SA6, which includes superior and transverse temporal cortices, closely recapitulates the auditory cortex. The functional specializations of the human auditory cortex^73^, which include parts of the lateral prefrontal cortex, Broca”s area, and subcentral regions, are needed for human vocalization and language^75,76^. The dorsal attention network (DAN), which directs voluntary allocation of attention, is concordant with SA1 that is comprised of superior parietal, supramarginal, postcentral, precuneus, isthmus cingulate, and inferior temporal regions. A noteworthy omission from SA1, which is an important feature of the DAN, are the frontal eye fields (FEF)^77^. The most likely explanation is that the FEF is not a distinct region in FreeSurfer parcellation and therefore this phenotype is not well represented in ENIGMA cortical GWAS. The DAN has relatively high twin heritability estimates (left h^2^=0.45; right h^2^=0.40)^74^. The default mode network (DMN), which is sometimes partitioned into dorsal and ventral subnetworks^78^, resembles SA2 and SA3 respectively. The caudal anterior cingulate, caudal middle frontal, medial orbitofrontal, paracentral, and rostral anterior cingulate cortices comprise SA2, while banks of STS, inferior parietal, and middle temporal cortices comprise SA3. SA4 represents a partial recapitulation of the central executive network (CEN) with the pars opercularis, pars orbitalis, and pars triangularis, but lacks the temporoparietal structures, which are a core feature of the CEN^79^.

Our results are consistent with evidence that the functional connectome is shaped by genomic constraints^74,80^. For instance, twin data from the human connectome project (HCP) shows that individual variability in the areal size of 17 canonical functional networks, as defined by Yeo et al.^73^, reveal marked heritability (h^2^=0.34 to 0.40). Unimodal sensory networks such as auditory and visual networks are particularly heritable while hetero-modal networks such as the salience network and the central executive network are significantly less heritable^74^. Overall, the parcellation schemes reflect the genetic influences driving cortical areal expansion and represent genetically driven processes in embryological neurodevelopment.

We find that behavioral traits and neuropsychiatric disorders showed distinct genetic correlations with SA-derived GIBNs that differ markedly from correlations with CT-derived GIBNs. Psychiatric disorders that were significantly genetically correlated with SA-derived GIBNS were not correlated with CT-derived GIBNs, and some CT-derived GIBNs were correlated with other psychiatric disorders that were not correlated with SA-derived GIBNs. CT3, which is located in the middle and superior temporal cortices, and CT4, which is located in the visual perceptual cortex, were strongly negatively correlated with alcohol use disorder. This divergent relationship between CT-derived and SA-derived networks is consistent with our findings form the ENIGMA-3 cortical GWAS where a consistent pattern of significant positive and negative correlations between total brain SA and behavioral traits/disorders was found, but average CT correlations with behavioral traits/disorders were non-significant^18^. Specifically, the ENIGMA-3 GWAS found that total SA was significantly positively correlated with cognitive function, educational attainment, Parkinson”s disease, and anorexia nervosa, but significantly negatively correlated with MDD, ADHD, depressive symptoms, neuroticism, and insomnia. In addition, the SA-derived GIBNs showed distinct genetic relationships to several psychiatric disorders. Several SA-derived GIBNs (SA1, SA2, SA4, SA5) were positively correlated with bipolar disorder, whereas SA-derived GIBNs (SA1, SA2, SA3, SA5, SA6) were negatively correlated with MDD, buttressing prior evidence that MDD and Bipolar are distinct conditions with divergent genetic bases^78^. While the relationship between these SA-derived GIBNs with MDD converge with the findings from the ENIGMA total surface area results, the relationship with bipolar disorder was novel. The GIBNs may provide additional power to detect genetic relationships when the strength of these relationships across cortical regions is heterogenous. Interestingly, although several GIBN-related SNPs we found were associated with schizophrenia, there were no GIBNs that were significantly correlated with schizophrenia (r_g_ =0.029 to 0.034; all p>0.30).

There is ample evidence that genetic variants that influence SA are distinct from genetic variants that influence CT^18^. Genetic variation affecting gene regulation in progenitor cell types, present in fetal development, affects adult cortical SA^81^. An increase in proliferative divisions of neural progenitor cells leads to an expansion of the pool of progenitors, resulting in increases in neuronal production and larger cortical SA, which is more prevalent in gyrencephalic species (e.g. humans, primates)^82^. By contrast, loci near genes implicated in cell differentiation, migration, adhesion, and myelination are associated with CT. The present findings suggest a similar distinction holds in case of SA-derived GIBNs compared to CT-derived GIBNs. We hypothesize that the unique genetic correlations of SA-derived GIBNs and CT-derived GIBNs with behavioral traits/disorders may be explained by the distinct developmental functions of their associated genes. Further exploration of the common and distinct relationships of SA-derived and CT-derived GIBNs with neuropsychiatric conditions using Mendelian Randomization and Latent Causal Variable (LCV) analysis may prove useful^83^.

### 4.1 Limitations

A number of limitations deserve consideration in interpreting the present findings. Our starting point for the gSEM was the GWAS of 34 cortical regions as defined by the Desikan-Killiany atlas. However, using cortical pleiotropy as an organizing schema for parcellation will not strictly adhere to regions defined by anatomical features. A high-resolution GWAS of the cortex would allow more flexibility in defining parcellation boundaries informed by genetic pleiotropy given it is likely that they differ from anatomically defined parcels. Realizing a high-resolution GWAS of the cortex poses a major computational challenge. For instance, a multivariate GWAS of 1,284 cortical vertex locations^26^ would be extremely time- and cost-prohibitive. Indeed, the present gSEM with just 34 phenotypes required about 2 weeks per chromosome running on the Shared Computing Cluster (SCC) at Boston University.

The GWAS threshold we used to identify significant associations with the 6 SA-derived and 4 CT-derived GIBNs was p<5×10^−8^, as we hypothesized that many of the associated loci would have been implicated in the prior GWAS, and we wanted to explore the relationship between GIBN GWAS variants and other traits. This is supported by the fact that the vast majority GIBN-associated SNPs had been noted in other cortical GWASs^17,18,29^. However, we note that the two novel SNPs identified, the SA1-associated SNP, rs3006933, would survive a strict Bonferroni correction for 10 GIBN GWASs examined (p=4.08×10^−9^), while the CT2-associated variant, rs1004763, would not (p=3.41×10^−08^). Therefore, the relevance of rs1004763 to cortical thickness should be considered provisional until replicated.

The present gSEM was based on the GWAS results of Grasby et al ^18^., which averaged left and right hemisphere phenotypic measures. This precluded an investigation of hemispheric asymmetries.

### 4.2 Conclusion

We harnessed the pervasive pleiotropy of the human cortex to realize a unique genetically-informed parcellation that is neurobiologically distinct from anatomical, functional, cytoarchitectural, and other established cortical parcellations, yet harbors meaningful topographic similarities to other network schemas, particularly resting-state fMRI networks. Strong genetic correlation between GIBNs and several major neuropsychiatric conditions including OCD, Bipolar, ADHD, and Alcohol Dependence, coupled with clear confirmation that nearly all GIBN-related SNPs play a role in cognitive, behavioral, neuroanatomical, and neurofunctional phenotypes, begins to expose the deeply interconnected architecture of the human cortex. Applying gSEM to model the joint genetic architecture of complex traits and investigate multivariate genetic links across phenotypes offers a new vantage point for mapping genetically informed cortical networks.

