## Supplementary Figures for "Genomic Structural Equation Modeling Reveals Latent Phenotypes in the Human Cortex with Distinct Genetic Architecture"

### Slide 1
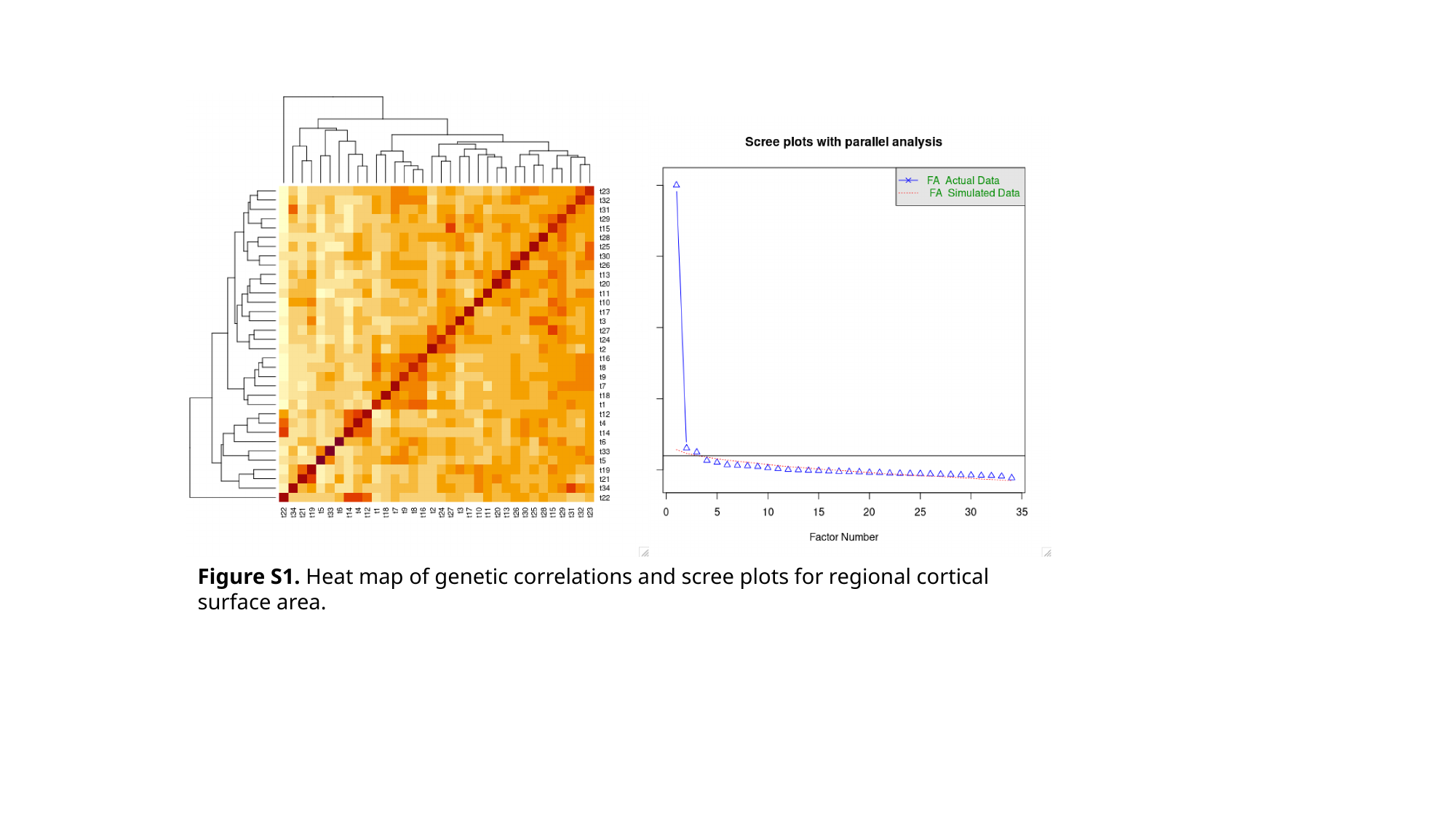

Figure S1. Heat map of genetic correlations and scree plots for regional cortical surface area.

### Slide 2
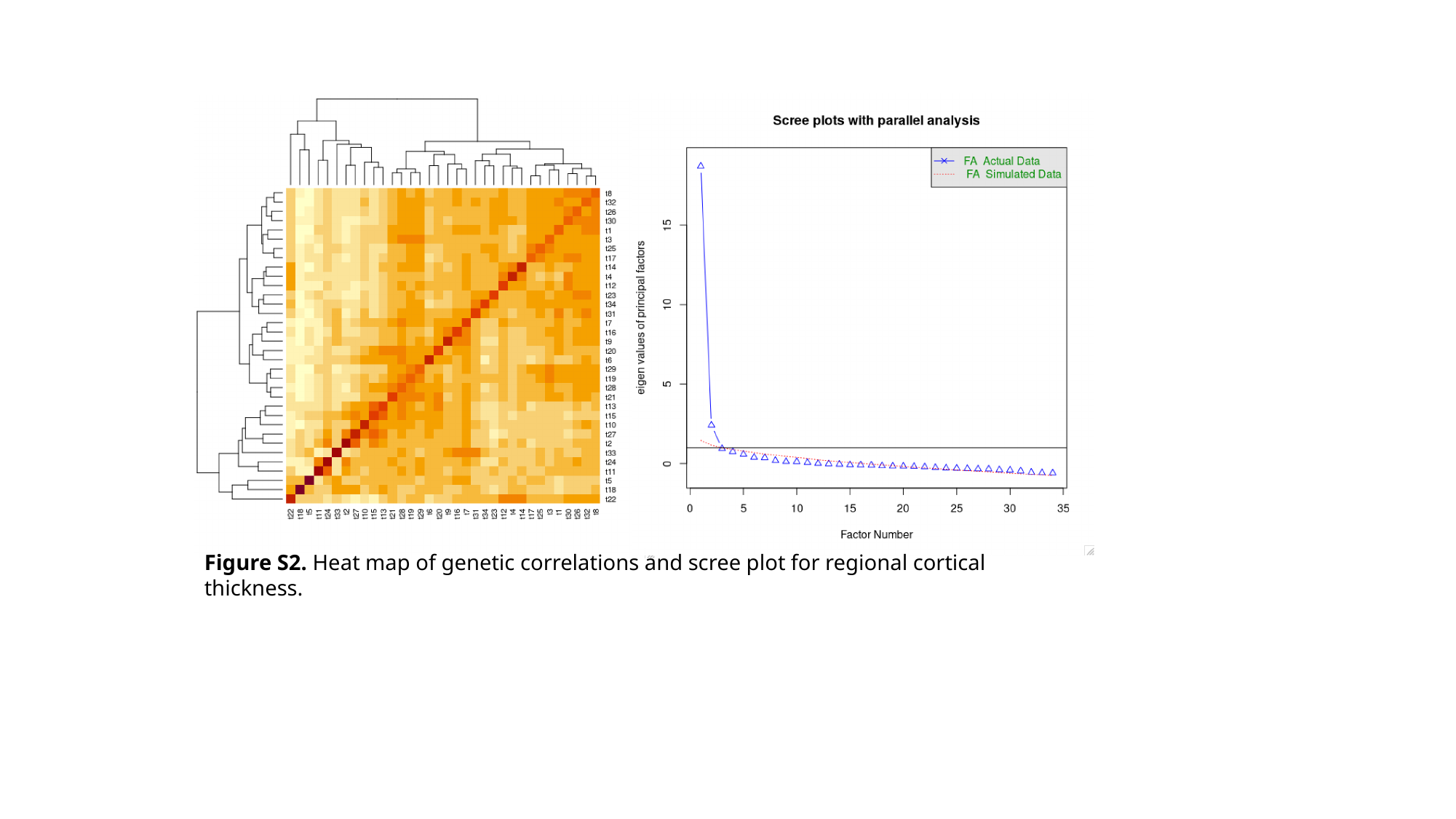

Figure S2. Heat map of genetic correlations and scree plot for regional cortical thickness.

### Slide 3
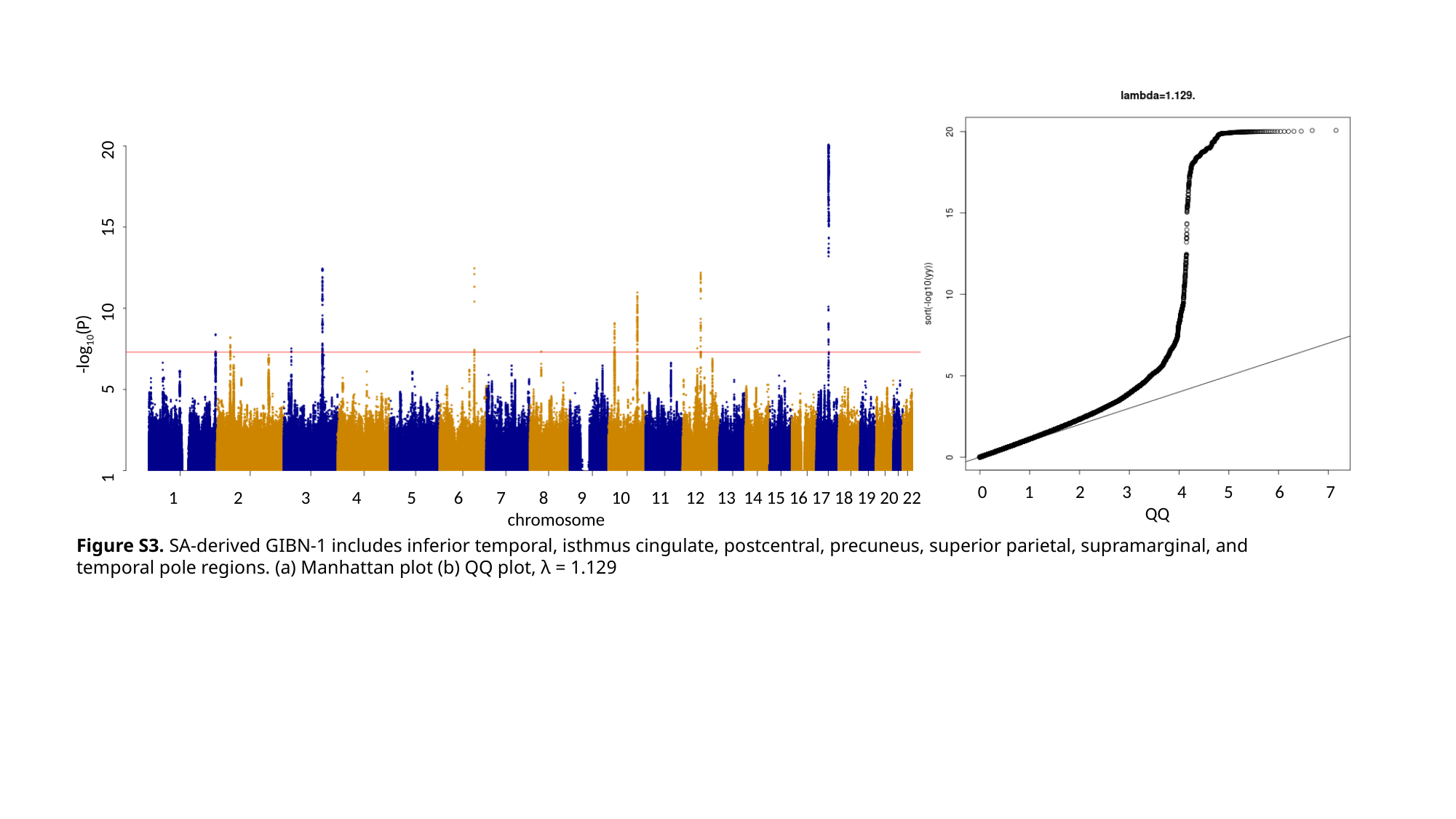

-log_10_(P)
1 5 10 15 20
0 1 2 3 4 5 6 7
 QQ
 2 3 4 5 6 7 8 9 10 11 12 13 14 15 16 17 18 19 20 22
 chromosome
Figure S3. SA-derived GIBN-1 includes inferior temporal, isthmus cingulate, postcentral, precuneus, superior parietal, supramarginal, and temporal pole regions. (a) Manhattan plot (b) QQ plot, λ = 1.129

### Slide 4
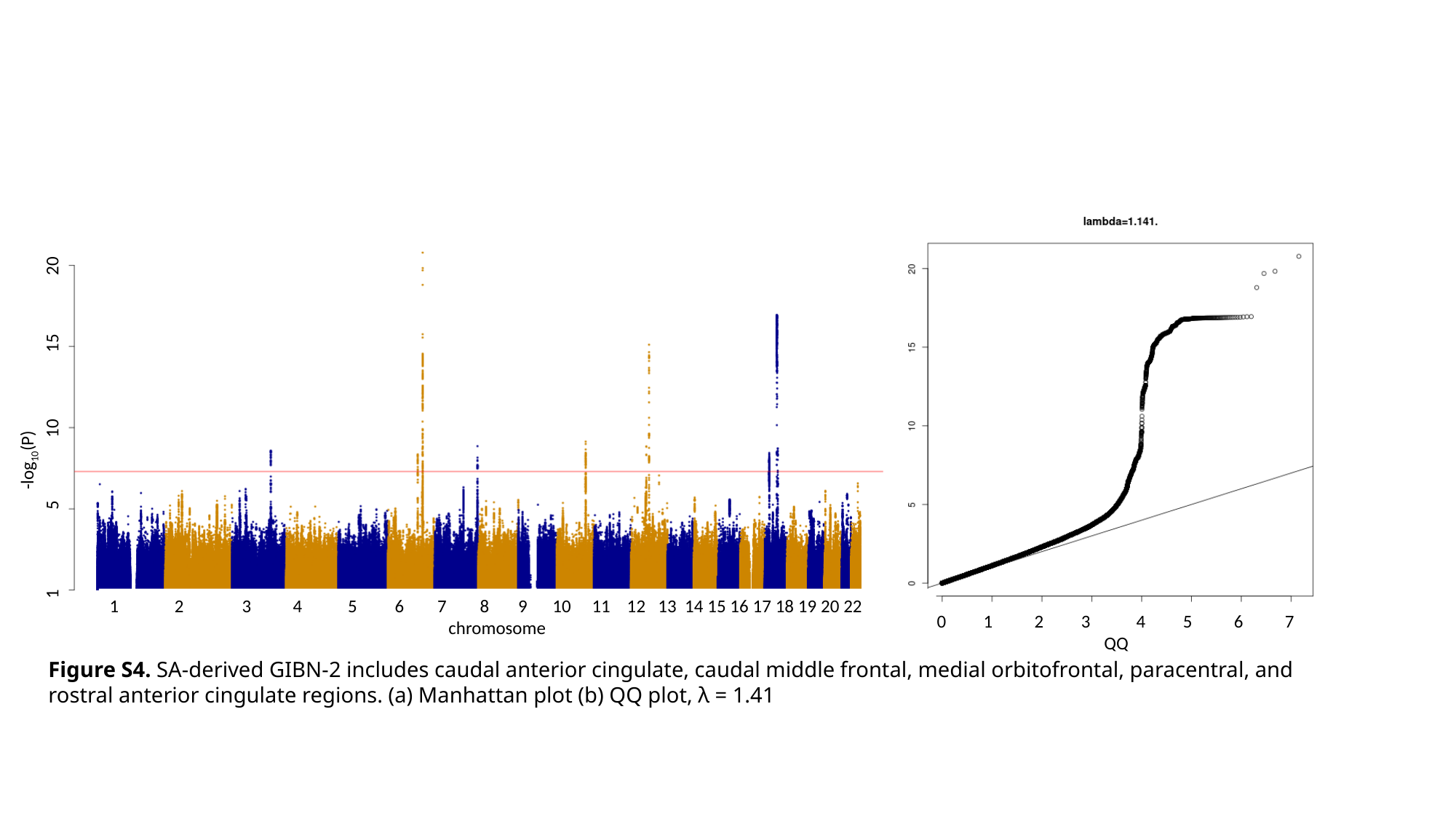

-log_10_(P)
1 5 10 15 20
 2 3 4 5 6 7 8 9 10 11 12 13 14 15 16 17 18 19 20 22
 chromosome
0 1 2 3 4 5 6 7
 QQ
Figure S4. SA-derived GIBN-2 includes caudal anterior cingulate, caudal middle frontal, medial orbitofrontal, paracentral, and rostral anterior cingulate regions. (a) Manhattan plot (b) QQ plot, λ = 1.41

### Slide 5
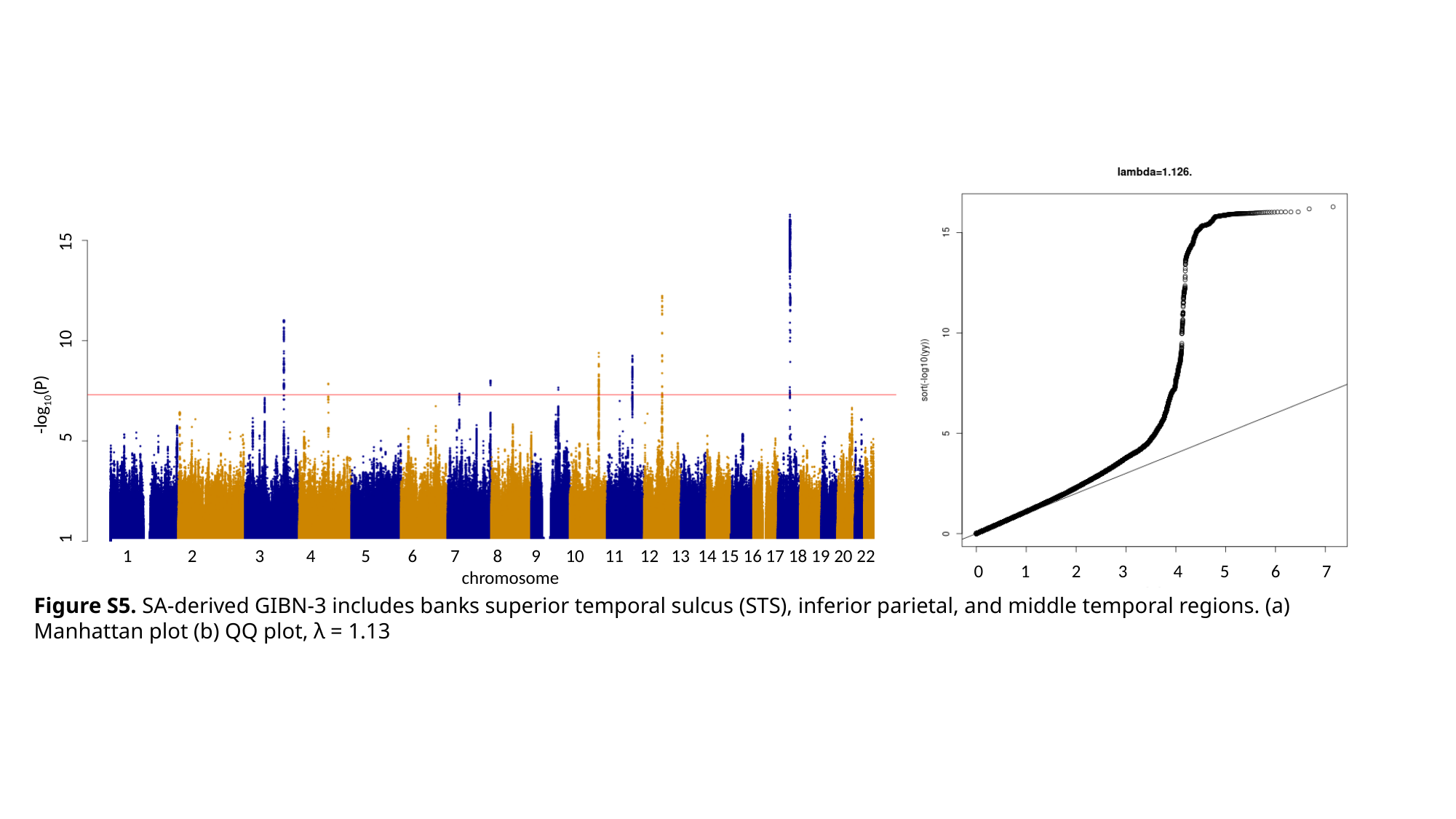

-log_10_(P)
1 5 10 15
 2 3 4 5 6 7 8 9 10 11 12 13 14 15 16 17 18 19 20 22
 chromosome
0 1 2 3 4 5 6 7
 QQ
Figure S5. SA-derived GIBN-3 includes banks superior temporal sulcus (STS), inferior parietal, and middle temporal regions. (a) Manhattan plot (b) QQ plot, λ = 1.13

### Slide 6
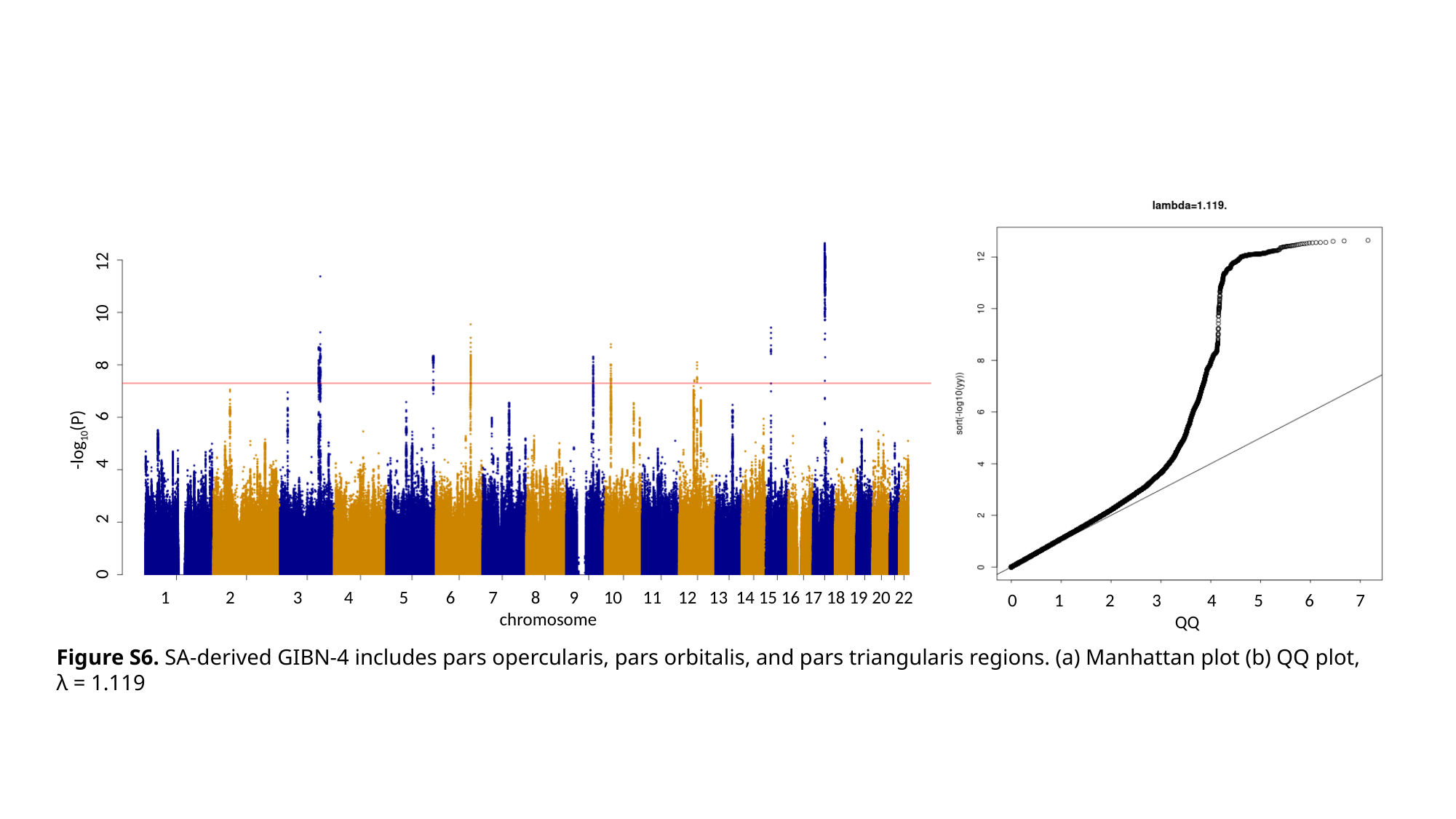

-log_10_(P)
0 2 4 6 8 10 12
 2 3 4 5 6 7 8 9 10 11 12 13 14 15 16 17 18 19 20 22
 chromosome
0 1 2 3 4 5 6 7
 QQ
Figure S6. SA-derived GIBN-4 includes pars opercularis, pars orbitalis, and pars triangularis regions. (a) Manhattan plot (b) QQ plot, λ = 1.119

### Slide 7
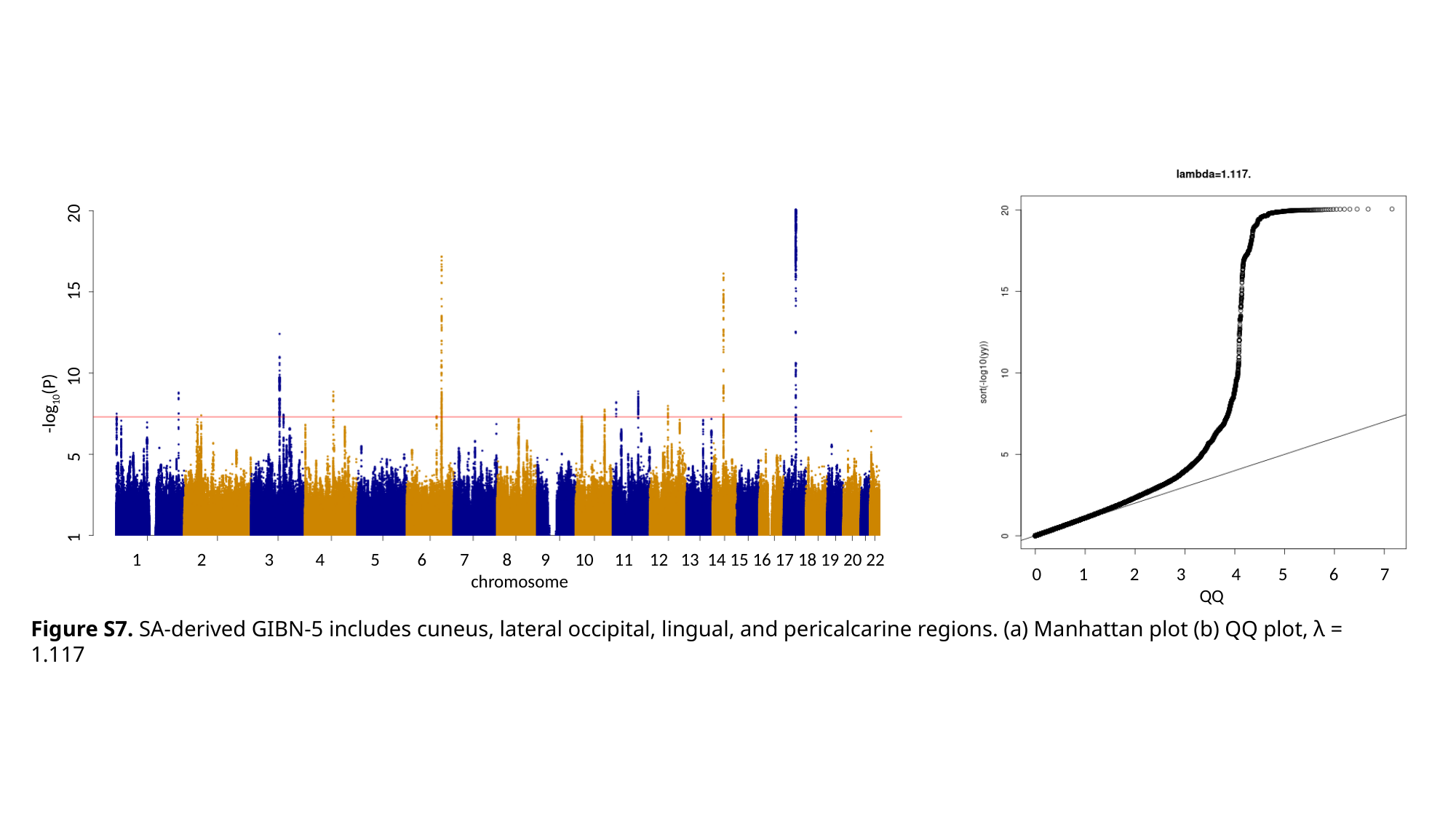

-log_10_(P)
1 5 10 15 20
 2 3 4 5 6 7 8 9 10 11 12 13 14 15 16 17 18 19 20 22
 chromosome
0 1 2 3 4 5 6 7
 QQ
Figure S7. SA-derived GIBN-5 includes cuneus, lateral occipital, lingual, and pericalcarine regions. (a) Manhattan plot (b) QQ plot, λ = 1.117

### Slide 8
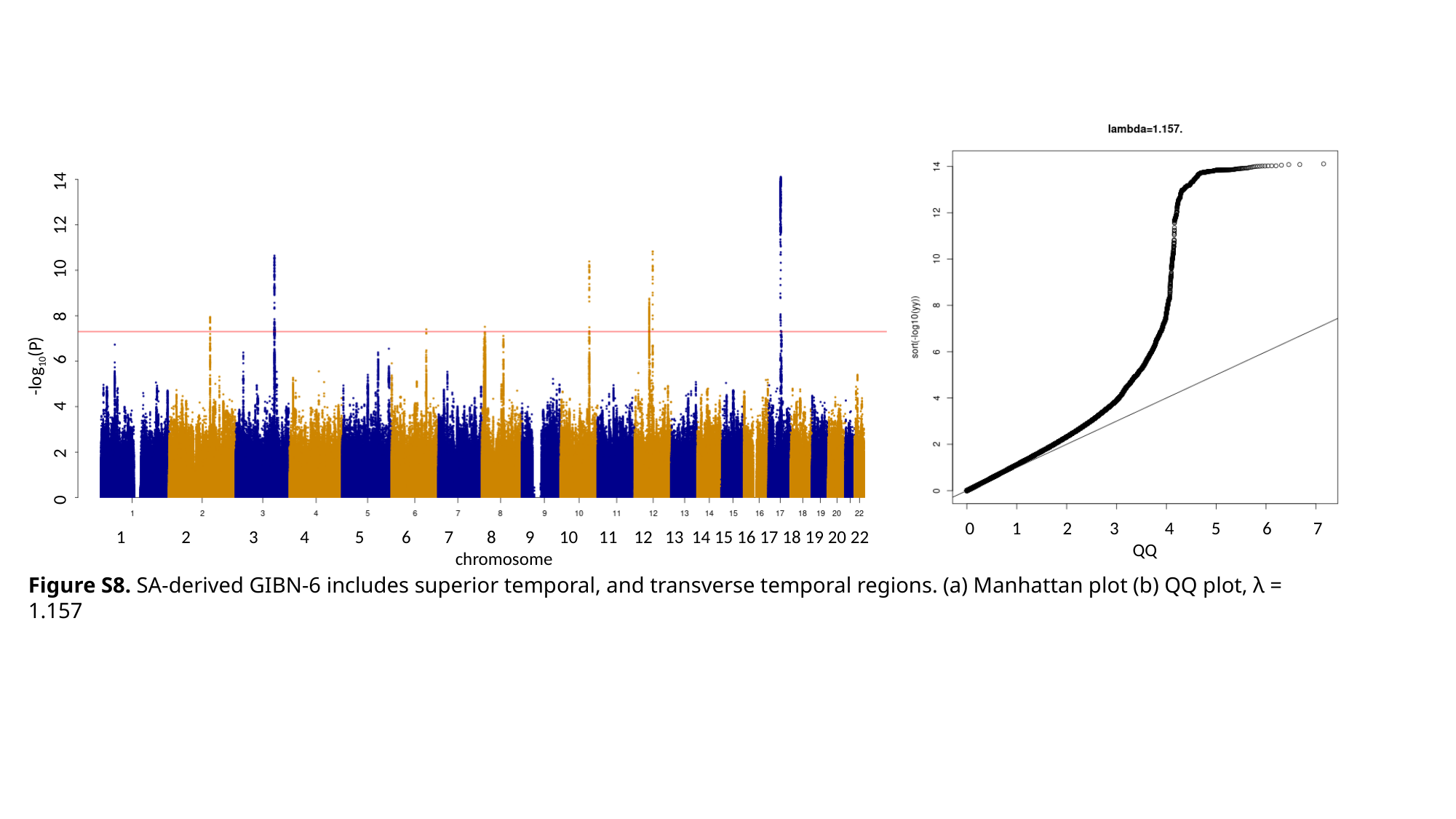

-log_10_(P)
0 2 4 6 8 10 12 14
0 1 2 3 4 5 6 7
 QQ
 2 3 4 5 6 7 8 9 10 11 12 13 14 15 16 17 18 19 20 22
 chromosome
Figure S8. SA-derived GIBN-6 includes superior temporal, and transverse temporal regions. (a) Manhattan plot (b) QQ plot, λ = 1.157

### Slide 9
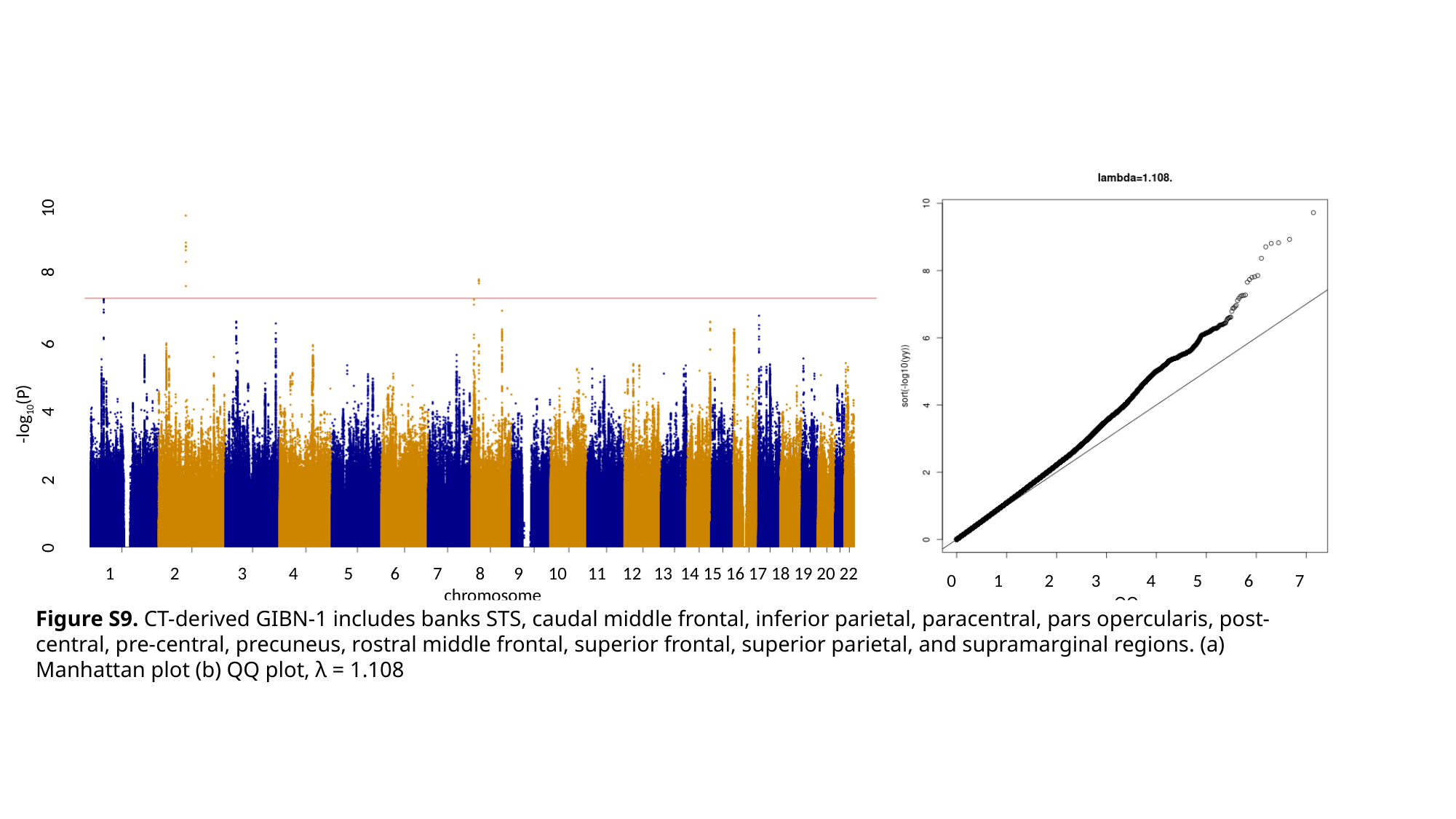

-log_10_(P)
0 2 4 6 8 10
 2 3 4 5 6 7 8 9 10 11 12 13 14 15 16 17 18 19 20 22
 chromosome
0 1 2 3 4 5 6 7
 QQ
Figure S9. CT-derived GIBN-1 includes banks STS, caudal middle frontal, inferior parietal, paracentral, pars opercularis, post-central, pre-central, precuneus, rostral middle frontal, superior frontal, superior parietal, and supramarginal regions. (a) Manhattan plot (b) QQ plot, λ = 1.108

### Slide 10
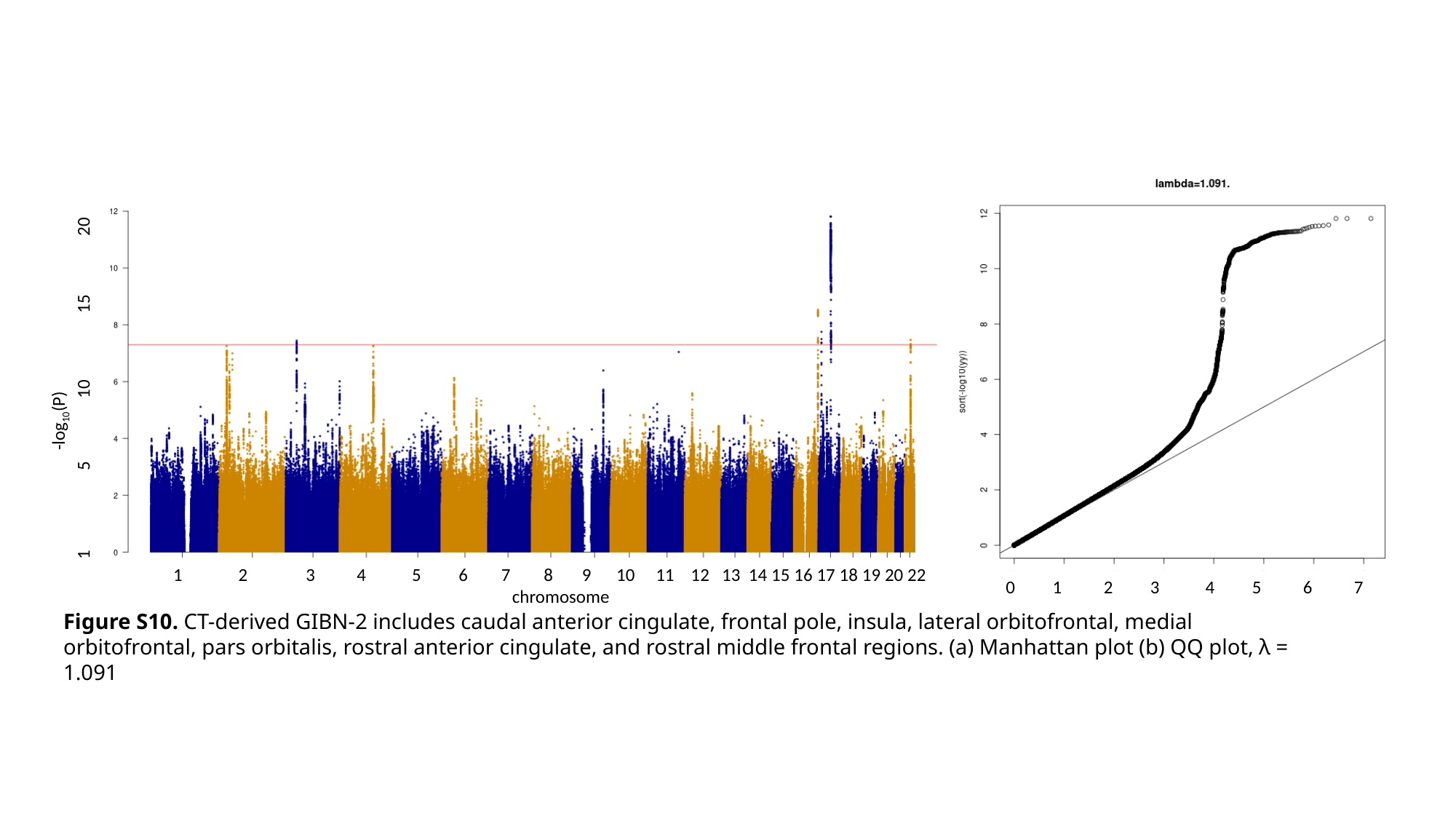

-log_10_(P)
1 5 10 15 20
 2 3 4 5 6 7 8 9 10 11 12 13 14 15 16 17 18 19 20 22
 chromosome
0 1 2 3 4 5 6 7
 QQ
Figure S10. CT-derived GIBN-2 includes caudal anterior cingulate, frontal pole, insula, lateral orbitofrontal, medial orbitofrontal, pars orbitalis, rostral anterior cingulate, and rostral middle frontal regions. (a) Manhattan plot (b) QQ plot, λ = 1.091

### Slide 11
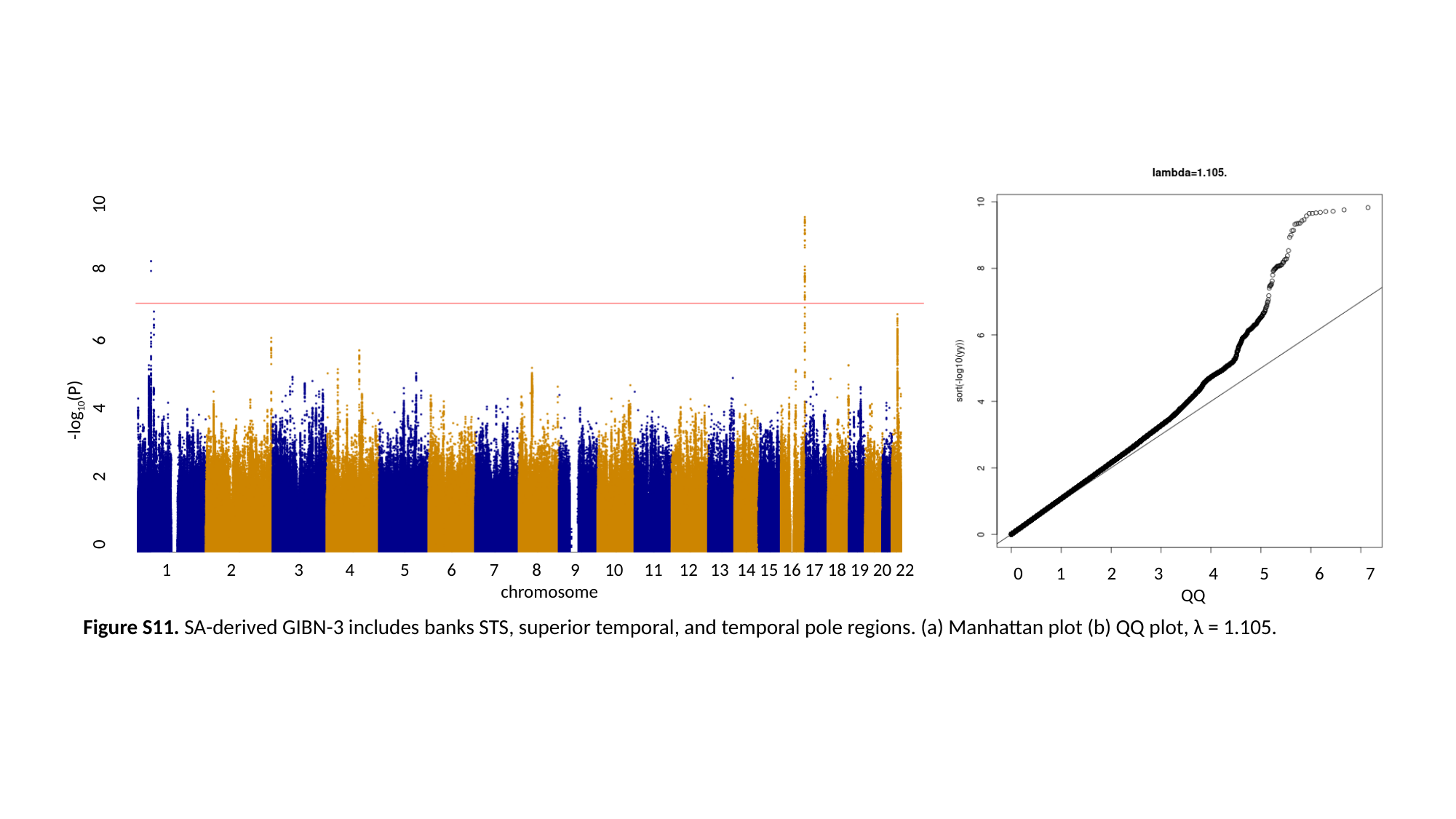

-log_10_(P)
0 2 4 6 8 10
 2 3 4 5 6 7 8 9 10 11 12 13 14 15 16 17 18 19 20 22
 chromosome
0 1 2 3 4 5 6 7
 QQ
Figure S11. SA-derived GIBN-3 includes banks STS, superior temporal, and temporal pole regions. (a) Manhattan plot (b) QQ plot, λ = 1.105.

### Slide 12
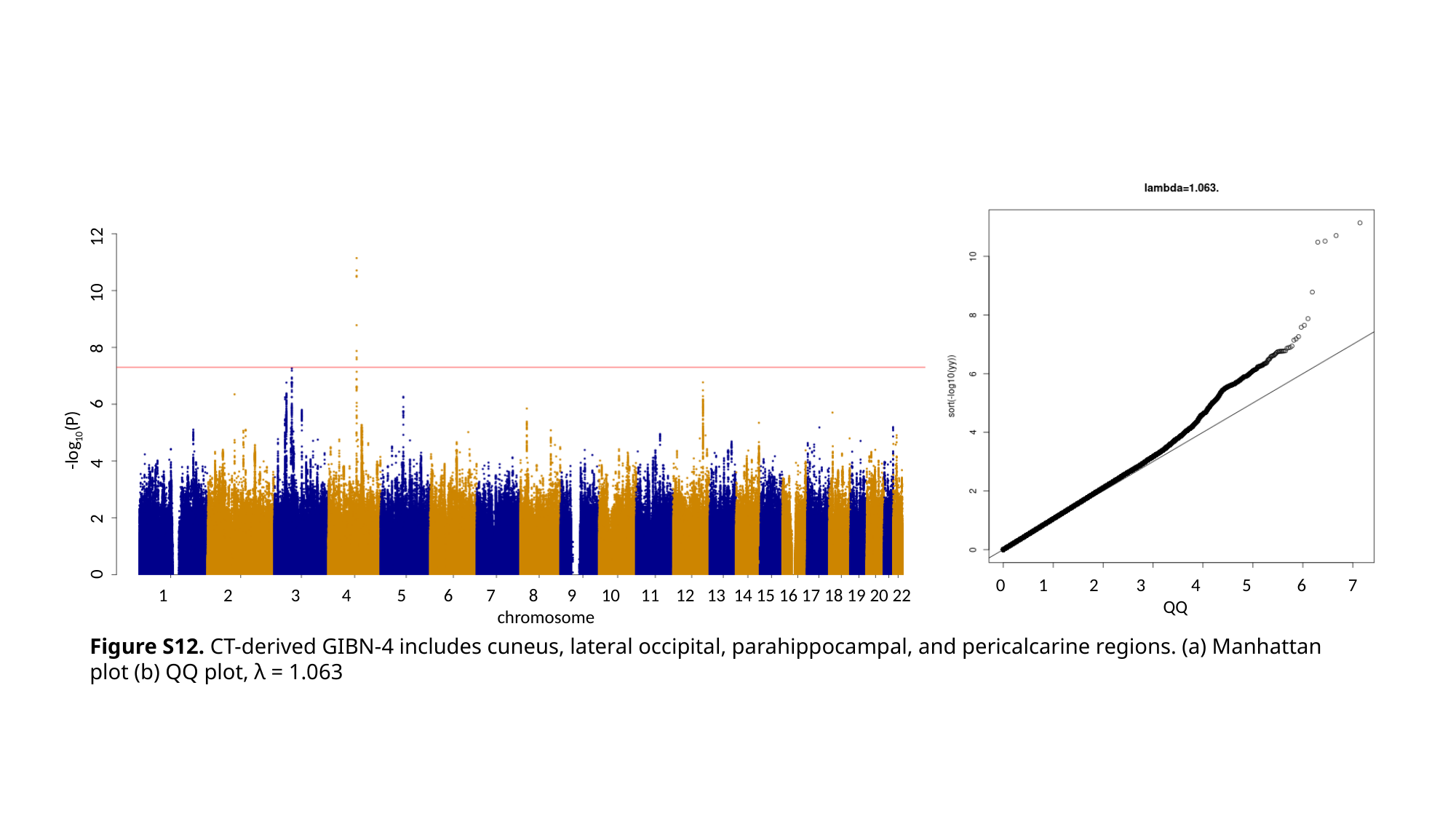

-log_10_(P)
0 2 4 6 8 10 12
0 1 2 3 4 5 6 7
 QQ
 2 3 4 5 6 7 8 9 10 11 12 13 14 15 16 17 18 19 20 22
 chromosome
Figure S12. CT-derived GIBN-4 includes cuneus, lateral occipital, parahippocampal, and pericalcarine regions. (a) Manhattan plot (b) QQ plot, λ = 1.063
