## Supplementary Tables for "Genomic Structural Equation Modeling Reveals Latent Phenotypes in the Human Cortex with Distinct Genetic Architecture"

**Table S1**. Model Fit Statistics for the genetically informed brain networks (GIBNS) of surface area. The 6 Factor surface area model fit the data the best (lower AIC, lower SRMR, or higher CFI indicate better fit).

| Number of Factors | Cutoff in EFA | AIC | model Ꭓ^2^ | DF | CFI | SRMR |
| --- | --- | --- | --- | --- | --- | --- |
| 6 | **0.5** | **22,712,604.05** | **22,712,478.1** | **237** | **0.92** | **0.062** |
| 5 | 0.5 | 30,128,114.56 | 30,127,992.6 | 264 | 0.91 | 0.064 |
| 3 | 0.5 | 37,107,176.74 | 37,107,062.7 | 321 | 0.89 | 0.064 |
| 6 | 0.3 | 38,521,514.25 | 38,521,322.3 | 465 | 0.91 | 0.056 |
| 9 | 0.3 | 39,497,504.82 | 39,497,276.8 | 481 | 0.91 | 0.055 |
| 4 | 0.3 | 42,768,842.21 | 42,768,674.2 | 511 | 0.90 | 0.058 |
| 7 | 0.3 | 43,175,172.83 | 43,174,968.8 | 493 | 0.90 | 0.060 |
| 5 | 0.3 | 44,590,135.39 | 44,589,955.4 | 471 | 0.90 | 0.059 |
| 3 | 0.3 | 47,348,969.86 | 47,348,817.9 | 519 | 0.89 | 0.061 |
| 2 | 0.5 | 51,228,607.05 | 51,228,473.1 | 494 | 0.88 | 0.071 |
| 1 | 0.5 | 53,156,095.52 | 53,155,963.5 | 495 | 0.87 | 0.073 |
| 2 | 0.3 | 53,561,525.01 | 53,561,385.0 | 525 | 0.88 | 0.071 |
| 1 | 0.3 | 60,204,948.95 | 60,204,812.9 | 527 | 0.86 | 0.077 |

Abbreviations: GIBN=genetically informed brain network; AIC=Akaike Information Criteria; CFI=Comparative Fit Index; SRMR=Standardized Root Mean Square Residuals.

**Table S2**. Estimated Loadings for the best fitting 6-GIBN model for surface area (standardized estimates).

| GIBN | Region | Standardized Estimates | p |
| --- | --- | --- | --- |
| SA1 | inferiortemporal | 0.79 | 6.25E-35 |
| SA1 | isthmuscingulate | 0.79 | 5.91E-25 |
| SA1 | postcentral | 0.90 | 8.90E-62 |
| SA1 | precuneus | 0.81 | 4.24E-44 |
| SA1 | superiorparietal | 0.82 | 8.00E-35 |
| SA1 | supramarginal | 0.90 | 5.45E-38 |
| SA1 | temporalpole | 0.70 | 6.73E-21 |
| SA2 | caudalanteriorcingulate | 0.76 | 7.75E-30 |
| SA2 | caudalmiddlefrontal | 0.76 | 4.78E-32 |
| SA2 | medialorbitofrontal | 0.93 | 1.36E-38 |
| SA2 | paracentral | 0.79 | 3.91E-31 |
| SA2 | rostralanteriorcingulate | 0.89 | 5.61E-48 |
| SA3 | bankssts | 0.87 | 4.12E-35 |
| SA3 | inferiorparietal | 0.91 | 1.04E-46 |
| SA3 | middletemporal | 0.96 | 3.97E-41 |
| SA4 | parsopercularis | 0.80 | 5.59E-25 |
| SA4 | parsorbitalis | 0.89 | 9.99E-32 |
| SA4 | parstriangularis | 0.68 | 1.69E-17 |
| SA5 | cuneus | 0.94 | 3.67E-51 |
| SA5 | lateraloccipital | 1.00 | 5.41E-56 |
| SA5 | lingual | 0.89 | 6.58E-39 |
| SA5 | pericalcarine | 0.66 | 1.86E-18 |
| SA6 | superiortemporal | 1.00 | 9.85E-35 |
| SA6 | transversetemporal | 0.83 | 3.96E-30 |

Abbreviations: GIBN=genetically informed brain network.

**Table S3**. Estimated correlation between GIBNS for the 6-GIBN model for surface area (standardized estimates).

| GIBN1 | GIBN2 | Correlation | p |
| --- | --- | --- | --- |
| SA1 | SA2 | 0.90 | < 5e-300 |
| SA1 | SA3 | 0.90 | 2.51E-213 |
| SA1 | SA4 | 0.84 | 1.16E-113 |
| SA1 | SA5 | 0.76 | 1.21E-94 |
| SA1 | SA6 | 0.78 | 1.70E-74 |
| SA2 | SA3 | 0.76 | 4.11E-79 |
| SA2 | SA4 | 0.91 | 3.77E-151 |
| SA2 | SA5 | 0.72 | 2.78E-74 |
| SA2 | SA6 | 0.80 | 1.67E-122 |
| SA3 | SA4 | 0.69 | 3.58E-39 |
| SA3 | SA5 | 0.61 | 1.29E-33 |
| SA3 | SA6 | 0.66 | 1.81E-27 |
| SA4 | SA5 | 0.73 | 1.32E-51 |
| SA4 | SA6 | 0.85 | 4.71E-49 |
| SA5 | SA6 | 0.64 | 1.62E-47 |

Abbreviations: GIBN=genetically informed brain network.

**Table S4**. Model Fit Statistics for genetically informed brain networks associated with cortical thickness. The 4-factor cortical thickness model fit the data the best (lower AIC and SRMR, higher CFI indicate better fit).

| Number of Factors | Cutoff in EFA | AIC | model Ꭓ^2^ | DF | CFI | SRMR |
| --- | --- | --- | --- | --- | --- | --- |
| 4 | **0.5** | **17,761,928** | **17,761,812.5** | **267** | **0.93** | **0.077** |
| 5 | 0.5 | 19,714,869 | 19,714,736.7 | 312 | 0.93 | 0.073 |
| 3 | 0.5 | 23,231,378 | 23,231,260.5 | 319 | 0.91 | 0.077 |
| 2 | 0.5 | 24,940,105 | 24,939,991.3 | 349 | 0.91 | 0.081 |
| 1 | 0.5 | 27,175,213 | 27,175,097.1 | 377 | 0.90 | 0.089 |
| 4 | 0.3 | 32,553,940 | 32,553,773.5 | 445 | 0.90 | 0.067 |
| 3 | 0.3 | 35,492,466 | 35,492,316.2 | 453 | 0.89 | 0.078 |
| 5 | 0.3 | 39,453,793 | 39,453,615.0 | 472 | 0.88 | 0.073 |
| 1 | 0.3 | 44,089,256 | 44,089,123.7 | 495 | 0.87 | 0.094 |

Abbreviations: AIC=Akaike Information Criteria; CFI=Comparative Fit Index; SRMR=Standardized Root Mean Square Residuals.

**Table S5**. Estimated loadings for the 4-GIBN model for cortical thickness (standardized estimates).

| GIBN | Region | standardized estimates | p |
| --- | --- | --- | --- |
| CT1 | bankssts | 0.83 | 1.74E-19 |
| CT1 | caudalmiddlefrontal | 0.87 | 8.69E-32 |
| CT1 | inferiorparietal | 0.92 | 3.80E-36 |
| CT1 | paracentral | 0.89 | 4.33E-44 |
| CT1 | parsopercularis | 0.90 | 8.81E-27 |
| CT1 | postcentral | 0.75 | 1.14E-15 |
| CT1 | precentral | 0.84 | 2.75E-24 |
| CT1 | precuneus | 0.95 | 1.67E-55 |
| CT1 | rostralmiddlefrontal | 0.54 | 1.37E-08 |
| CT1 | superiorfrontal | 0.93 | 9.93E-39 |
| CT1 | superiorparietal | 0.89 | 4.56E-39 |
| CT1 | supramarginal | 0.92 | 1.38E-30 |
| CT2 | caudalanteriorcingulate | 0.65 | 4.17E-12 |
| CT2 | frontalpole | 0.86 | 1.39E-13 |
| CT2 | insula | 0.81 | 8.09E-20 |
| CT2 | lateralorbitofrontal | 0.87 | 1.52E-20 |
| CT2 | medialorbitofrontal | 0.74 | 1.62E-17 |
| CT2 | parsorbitalis | 0.92 | 2.98E-21 |
| CT2 | rostralanteriorcingulate | 0.67 | 2.42E-10 |
| CT2 | rostralmiddlefrontal | 0.43 | 2.37E-05 |
| CT3 | superiortemporal | 0.98 | 2.10E-32 |
| CT3 | temporalpole | 0.71 | 1.58E-09 |
| CT4 | cuneus | 0.76 | 2.52E-18 |
| CT4 | lateraloccipital | 0.92 | 1.27E-31 |
| CT4 | parahippocampal | 0.40 | 2.55E-06 |
| CT4 | pericalcarine | 0.66 | 2.86E-09 |

Abbreviations: GIBN=genetically informed brain network;

**Table S6**. Estimated correlation between GIBNS for the 6-GIBN model for cortical thickness as standardized estimates (ρ).

| GIBN1 | GIBN2 | ρ | p |
| --- | --- | --- | --- |
| CT1 | CT2 | 0.71 | 2.57E-47 |
| CT1 | CT3 | 0.77 | 4.42E-31 |
| CT1 | CT4 | 0.87 | 2.13E-53 |
| CT2 | CT3 | 0.76 | 8.02E-28 |
| CT2 | CT4 | 0.67 | 1.10E-19 |
| CT3 | CT4 | 0.68 | 3.68E-17 |

Abbreviations: GIBN=genetically informed brain network

**Table S7**. All Genome-wide significant variants associated with GINBs

| **GIBN** | **rsID** | **Chr** | **Pos** | **Beta** | **p** |
| --- | --- | --- | --- | --- | --- |
| **SA2** | rs4273712 | 6 | 126964510 | -0.29 | 1.66E-21 |
| **SA1** | rs62057153 | 17 | 43904528 | 0.28 | 8.45E-21 |
| **SA5** | rs55638417 | 17 | 43900434 | 0.28 | 8.60E-21 |
| **SA5** | rs74580701 | 6 | 127000881 | 0.66 | 6.64E-18 |
| **SA2** | rs79600142 | 17 | 43897722 | 0.26 | 1.13E-17 |
| **SA3** | rs56319902 | 17 | 43871982 | 0.26 | 5.22E-17 |
| **SA5** | rs73313052 | 14 | 59625997 | 0.28 | 7.48E-17 |
| **SA2** | rs7312464 | 12 | 66374247 | 0.20 | 7.78E-16 |
| **SA6** | rs9896243 | 17 | 44826056 | 0.23 | 7.75E-15 |
| **SA4** | rs55663797 | 17 | 43544379 | 0.27 | 2.27E-13 |
| **SA1** | rs11759026 | 6 | 126792095 | -0.19 | 3.48E-13 |
| **SA1** | rs34464850 | 3 | 141721762 | -0.21 | 3.60E-13 |
| **SA5** | rs6788676 | 3 | 104674040 | -0.17 | 3.89E-13 |
| **SA3** | rs8756 | 12 | 66359752 | 0.18 | 5.84E-13 |
| **SA1** | rs7312464 | 12 | 66374247 | 0.16 | 6.47E-13 |
| **CT2** | rs2316766 | 17 | 43919068 | -0.28 | 1.54E-12 |
| **SA4** | rs2279829 | 3 | 147106319 | -0.24 | 4.21E-12 |
| **CT4** | rs13107325 | 4 | 103188709 | 0.44 | 7.19E-12 |
| **SA3** | rs34464850 | 3 | 141721762 | -0.21 | 9.61E-12 |
| **SA1** | rs1628768 | 10 | 105012994 | -0.17 | 1.07E-11 |
| **SA6** | rs10878349 | 12 | 66327632 | 0.16 | 1.49E-11 |
| **SA6** | rs34464850 | 3 | 141721762 | -0.20 | 2.27E-11 |
| **SA6** | rs1163249 | 10 | 104909890 | 0.18 | 4.10E-11 |
| **CT3** | rs12711473 | 16 | 87224293 | 0.19 | 1.48E-10 |
| **CT1** | rs11692435 | 2 | 98275354 | 0.30 | 1.90E-10 |
| **SA4** | rs11759026 | 6 | 126792095 | -0.19 | 2.84E-10 |
| **SA4** | rs4924345 | 15 | 39639898 | 0.32 | 3.78E-10 |
| **SA3** | rs4917384 | 10 | 104995788 | 0.15 | 4.13E-10 |
| **SA3** | rs3847535 | 11 | 92558024 | 0.15 | 5.62E-10 |
| **SA2** | rs1628768 | 10 | 105012994 | -0.16 | 7.21E-10 |
| **SA1** | rs7082934 | 10 | 21881379 | 0.13 | 8.26E-10 |
| **SA5** | rs3858368 | 11 | 92553286 | 0.14 | 1.35E-09 |
| **SA2** | rs7778997 | 7 | 156183533 | -0.36 | 1.38E-09 |
| **SA5** | rs13135092 | 4 | 103198082 | -0.23 | 1.41E-09 |
| **SA2** | rs7297175 | 12 | 56473808 | 0.14 | 1.43E-09 |
| **SA5** | rs3754356 | 1 | 228270964 | 0.16 | 1.61E-09 |
| **SA4** | rs11012732 | 10 | 21830104 | 0.17 | 1.63E-09 |
| **SA6** | rs11170566 | 12 | 53907067 | -0.19 | 1.78E-09 |
| **SA2** | rs11079849 | 17 | 47090785 | -0.15 | 1.91E-09 |
| **SA4** | rs2271386 | 3 | 141712708 | -0.20 | 2.12E-09 |
| **SA2** | rs3817176 | 3 | 141712780 | -0.17 | 2.64E-09 |
| **CT3** | rs61784835 | 1 | 47974123 | 0.17 | 2.92E-09 |
| **CT2** | rs12711473 | 16 | 87224293 | 0.17 | 2.94E-09 |
| **SA2** | rs4792721 | 17 | 16006827 | 0.13 | 3.60E-09 |
| **SA1** | rs3006933 | 1 | 243659727 | -0.13 | 4.08E-09 |
| **SA2** | rs2490272 | 6 | 108895386 | -0.13 | 4.41E-09 |
| **SA4** | rs7711765 | 5 | 170832637 | -0.16 | 4.52E-09 |
| **SA4** | rs113343136 | 9 | 98314306 | -0.26 | 4.88E-09 |
| **SA1** | rs7563432 | 2 | 48281138 | 0.13 | 6.18E-09 |
| **SA5** | rs10765918 | 11 | 12071855 | 0.15 | 6.24E-09 |
| **SA4** | rs7312464 | 12 | 66374247 | 0.15 | 7.94E-09 |
| **SA3** | rs78200999 | 7 | 155996837 | -0.14 | 9.78E-09 |
| **SA5** | rs10878349 | 12 | 66327632 | 0.12 | 1.06E-08 |
| **SA6** | rs13035861 | 2 | 150040224 | -0.12 | 1.13E-08 |
| **SA3** | rs2301718 | 4 | 106009763 | -0.16 | 1.39E-08 |
| **CT1** | rs3200031 | 8 | 26227484 | -0.24 | 1.41E-08 |
| **CT2** | rs12602519 | 17 | 10027480 | -0.16 | 1.77E-08 |
| **SA5** | rs1628768 | 10 | 105012994 | -0.13 | 1.77E-08 |
| **SA3** | rs28457693 | 9 | 98217348 | -0.20 | 2.18E-08 |
| **SA1** | rs7975351 | 12 | 53902190 | -0.12 | 2.93E-08 |
| **SA1** | rs12630663 | 3 | 28007315 | -0.12 | 3.02E-08 |
| **SA6** | rs9969436 | 8 | 10842659 | -0.12 | 3.07E-08 |
| **SA5** | rs6701689 | 1 | 2020343 | -0.13 | 3.19E-08 |
| **CT2** | rs1004763 | 22 | 38474952 | -0.15 | 3.41E-08 |
| **SA5** | rs56007616 | 3 | 118994959 | 0.18 | 3.61E-08 |
| **CT2** | rs533577 | 3 | 39489651 | 0.15 | 3.61E-08 |
| **SA4** | rs7297175 | 12 | 56473808 | 0.14 | 3.83E-08 |
| **SA6** | rs4273712 | 6 | 126964510 | -0.13 | 3.98E-08 |
| **SA5** | rs7570830 | 2 | 61745528 | -0.11 | 4.01E-08 |
| **SA3** | rs2299148 | 7 | 42023400 | -0.14 | 4.43E-08 |
| **SA5** | rs2802292 | 6 | 108908518 | -0.11 | 4.63E-08 |
| **SA1** | rs10109434 | 8 | 41169496 | 0.12 | 4.72E-08 |
| **SA5** | rs12357321 | 10 | 21790476 | 0.12 | 4.74E-08 |
| **SA6** | rs9909861 | 17 | 47079416 | -0.12 | 4.89E-08 |

**Table S8**. LDSC estimates of the pairwise genetic correlation between SA-derived GIBNs and psychiatric disorders.

| GIBN | Disorder | r_g_ | p | p_FDR_ |
| --- | --- | --- | --- | --- |
| SA3 | ADHD | -0.20 | **3.29E-06** | **0.00040** |
| SA4 | Bipolar | 0.14 | **3.00E-04** | **0.012** |
| SA4 | Cannabis | 0.15 | **4.00E-04** | **0.012** |
| SA1 | ADHD | -0.15 | **6.00E-04** | **0.014** |
| SA6 | ADHD | -0.16 | **8.00E-04** | **0.014** |
| SA3 | MDD | -0.10 | **0.0011** | **0.017** |
| SA5 | ADHD | -0.13 | **0.0013** | **0.017** |
| SA2 | Bipolar | 0.13 | **0.0018** | **0.022** |
| SA2 | ADHD | -0.13 | **0.0038** | **0.039** |
| SA5 | Bipolar | 0.10 | **0.0039** | **0.039** |
| SA1 | Bipolar | 0.11 | **0.0047** | **0.043** |
| SA1 | MDD | -0.080 | **0.0098** | 0.078 |
| SA2 | Cannabis | 0.11 | **0.011** | 0.080 |
| SA2 | MDD | -0.075 | **0.020** | 0.14 |
| SA6 | MDD | -0.071 | **0.030** | 0.20 |
| SA6 | Cannabis | 0.082 | **0.039** | 0.23 |
| SA5 | MDD | -0.057 | **0.046** | 0.26 |
| SA3 | PTSD | -0.089 | 0.056 | 0.29 |
| SA3 | Cannabis | 0.075 | 0.085 | 0.37 |
| SA1 | Tourette’s | -0.099 | 0.087 | 0.37 |
| SA4 | ADHD | -0.076 | 0.093 | 0.39 |
| SA2 | Autism | 0.086 | 0.114 | 0.40 |
| SA6 | Autism | 0.11 | 0.124 | 0.41 |
| SA4 | Autism | 0.080 | 0.125 | 0.41 |
| SA3 | Bipolar | 0.061 | 0.131 | 0.41 |
| SA5 | Tourette’s | -0.086 | 0.142 | 0.44 |
| SA4 | MDD | -0.044 | 0.152 | 0.45 |
| SA6 | Tourette’s | -0.088 | 0.153 | 0.45 |
| SA4 | Anxiety | 0.17 | 0.18 | 0.49 |
| SA2 | Tourette’s | -0.090 | 0.19 | 0.51 |
| SA2 | Alcohol Dependence | 0.097 | 0.21 | 0.54 |
| SA6 | Bipolar | 0.047 | 0.21 | 0.54 |
| SA1 | Autism | 0.058 | 0.23 | 0.57 |
| SA2 | PTSD | -0.054 | 0.23 | 0.57 |
| SA3 | Tourette’s | -0.079 | 0.24 | 0.57 |
| SA1 | Cannabis | 0.049 | 0.24 | 0.57 |
| SA1 | Alcohol Dependence | 0.090 | 0.25 | 0.58 |
| SA6 | OCD | 0.079 | 0.27 | 0.58 |
| SA2 | OCD | 0.078 | 0.27 | 0.58 |
| SA3 | Anorexia Nervosa | -0.047 | 0.27 | 0.58 |
| SA3 | Schizophrenia | -0.029 | 0.35 | 0.71 |
| SA5 | Anorexia Nervosa | 0.034 | 0.38 | 0.71 |
| SA3 | OCD | 0.064 | 0.38 | 0.71 |
| SA6 | Schizophrenia | -0.026 | 0.39 | 0.71 |
| SA5 | PTSD | -0.038 | 0.39 | 0.71 |
| SA6 | PTSD | -0.033 | 0.45 | 0.77 |
| SA1 | OCD | 0.051 | 0.46 | 0.77 |
| SA3 | Anxiety | -0.070 | 0.50 | 0.82 |
| SA4 | OCD | 0.046 | 0.55 | 0.85 |
| SA5 | Autism | 0.031 | 0.55 | 0.85 |
| SA2 | Anxiety | 0.065 | 0.58 | 0.88 |
| SA6 | Anorexia Nervosa | -0.023 | 0.59 | 0.88 |
| SA1 | PTSD | -0.025 | 0.59 | 0.88 |
| SA4 | PTSD | -0.022 | 0.64 | 0.91 |
| SA2 | Schizophrenia | 0.014 | 0.66 | 0.91 |
| SA5 | Schizophrenia | -0.013 | 0.67 | 0.91 |
| SA5 | Alcohol Dependence | -0.029 | 0.69 | 0.91 |
| SA3 | Alcohol Dependence | 0.031 | 0.70 | 0.91 |
| SA1 | Schizophrenia | -0.011 | 0.73 | 0.93 |
| SA4 | Tourette’s | 0.021 | 0.74 | 0.94 |
| SA4 | Alcohol Dependence | 0.026 | 0.75 | 0.94 |
| SA2 | Anorexia Nervosa | -0.012 | 0.78 | 0.95 |
| SA5 | Anxiety | 0.028 | 0.78 | 0.95 |
| SA1 | Anorexia Nervosa | -0.011 | 0.79 | 0.95 |
| SA6 | Anxiety | -0.034 | 0.80 | 0.95 |
| SA4 | Anorexia Nervosa | 0.012 | 0.81 | 0.95 |
| SA5 | Cannabis | -0.0089 | 0.81 | 0.95 |
| SA3 | Autism | -0.0084 | 0.87 | 0.96 |
| SA1 | Anxiety | -0.017 | 0.88 | 0.96 |
| SA6 | Alcohol Dependence | -0.010 | 0.89 | 0.97 |
| SA5 | OCD | -0.0073 | 0.92 | 0.99 |
| SA4 | Schizophrenia | 0.0015 | 0.96 | 0.99 |

Abbreviations: LDSC=LD score regression; GIBN=genetically informed brain network; SA- =surface area derived GIBNS; r_g_=genetic correlation; ADHD=Attention Deficit Hyperactivity Disorder; MDD=Major Depressive Disorder; P_FDR_ False Discovery Rate corrected significance value.

**Table S9**. LDSC estimates of the pairwise genetic correlation between CT-derived GIBNs and psychiatric disorders. Nominally significant (p<0.05) associations presented.

| GIBN | Disorder | r_g_ | p | p_FDR_ |
| --- | --- | --- | --- | --- |
| CT3 | Alcohol Dependence | -0.35 | **3.00E-04** | **0.012** |
| CT2 | Alcohol Dependence | -0.31 | **7.00E-04** | **0.014** |
| CT3 | OCD | 0.22 | **0.0091** | 0.078 |
| CT1 | Alcohol Dependence | -0.18 | **0.035** | 0.22 |
| CT3 | Anxiety | -0.27 | 0.051 | 0.28 |
| CT1 | Anorexia Nervosa | -0.078 | 0.060 | 0.30 |
| CT4 | Alcohol Dependence | -0.16 | 0.084 | 0.37 |
| CT1 | Anxiety | -0.21 | 0.087 | 0.37 |
| CT2 | Cannabis | -0.079 | 0.10 | 0.40 |
| CT4 | Anxiety | -0.20 | 0.11 | 0.40 |
| CT1 | OCD | 0.13 | 0.11 | 0.40 |
| CT1 | Tourettes | 0.11 | 0.11 | 0.40 |
| CT4 | OCD | 0.12 | 0.13 | 0.41 |
| CT2 | Anorexia Nervosa | -0.063 | 0.17 | 0.49 |
| CT2 | PTSD | -0.072 | 0.17 | 0.49 |
| CT3 | Tourette’s | 0.077 | 0.26 | 0.58 |
| CT1 | Bipolar | 0.042 | 0.31 | 0.64 |
| CT4 | Autism | 0.056 | 0.31 | 0.64 |
| CT4 | Schizophrenia | 0.033 | 0.34 | 0.69 |
| CT4 | Cannabis | -0.044 | 0.37 | 0.71 |
| CT2 | ADHD | -0.043 | 0.38 | 0.71 |
| CT1 | PTSD | -0.037 | 0.43 | 0.77 |
| CT4 | Bipolar | 0.035 | 0.44 | 0.77 |
| CT3 | PTSD | -0.037 | 0.45 | 0.77 |
| CT3 | Cannabis | -0.040 | 0.46 | 0.77 |
| CT4 | ADHD | 0.032 | 0.51 | 0.83 |
| CT1 | Autism | -0.047 | 0.53 | 0.84 |
| CT3 | MDD | -0.021 | 0.59 | 0.88 |
| CT1 | MDD | 0.018 | 0.61 | 0.89 |
| CT3 | Schizophrenia | 0.017 | 0.61 | 0.89 |
| CT4 | Anorexia Nervosa | -0.022 | 0.63 | 0.90 |
| CT3 | Bipolar | 0.020 | 0.66 | 0.91 |
| CT3 | Autism | -0.032 | 0.69 | 0.91 |
| CT4 | Tourette’s | 0.027 | 0.70 | 0.91 |
| CT1 | Schizophrenia | 0.012 | 0.71 | 0.91 |
| CT2 | Schizophrenia | 0.011 | 0.76 | 0.94 |
| CT2 | Autism | -0.013 | 0.86 | 0.96 |
| CT3 | ADHD | -0.0086 | 0.87 | 0.96 |
| CT2 | OCD | -0.014 | 0.87 | 0.96 |
| CT3 | Anorexia Nervosa | -0.0072 | 0.88 | 0.96 |
| CT2 | Bipolar | -0.0044 | 0.93 | 0.99 |
| CT1 | Cannabis | 0.004 | 0.94 | 0.99 |
| CT1 | ADHD | -0.0032 | 0.95 | 0.99 |
| CT2 | Anxiety | -0.0050 | 0.97 | 0.99 |
| CT4 | PTSD | 0.0012 | 0.98 | 0.99 |
| CT4 | MDD | -0.00060 | 0.99 | 0.99 |
| CT2 | Tourette’s | -0.0010 | 0.99 | 0.99 |
| CT2 | MDD | 0.00030 | 0.99 | 0.99 |

Abbreviations: GIBN=genetically informed brain network; CT- =cortical thickness derived GIBNS; r_g_=genetic correlation; ADHD=Attention Deficit Hyperactivity Disorder; MDD=Major Depressive Disorder; P_FDR_ False Discovery Rate corrected significance value.
